# Inferring the landscapes of mutation and recombination in the common marmoset (*Callithrix jacchus*) in the presence of twinning and hematopoietic chimerism

**DOI:** 10.1101/2025.07.01.662565

**Authors:** Vivak Soni, Cyril J. Versoza, Jeffrey D. Jensen, Susanne P. Pfeifer

## Abstract

The common marmoset (*Callithrix jacchus*) is an important model in biomedical and clinical research, particularly for the study of age-related, neurodegenerative, and neurodevelopmental disorders (due to their biological similarities with humans), infectious disease (due to their susceptibility to a variety of pathogens), as well as developmental biology (due to their short gestation period relative to many other primates). Yet, while being one of the most commonly used non-human primate models for research, the population genomics of the common marmoset remains relatively poorly characterized, despite the critical importance of this knowledge in many areas of research including genome-wide association studies, models of polygenic risk scores, and scans for the targets of selection. This neglect owes, at least in part, to two biological peculiarities related to the reproductive mode of the species — frequent twinning and sibling chimerism — which are likely to affect standard population genetic approaches relying on assumptions underlying the Wright-Fisher model. Using high-quality population genomic data, we here infer the rates and landscapes of mutation and recombination — two fundamental processes dictating the levels and patterns of genetic variability — in the presence of these biological features, and discuss our findings in light of recent work in primates. Our results suggest that, while the species exhibits relatively low neutral mutation rates, rates of recombination are in the range of those observed in other anthropoids. Moreover, the recombination landscape of common marmosets, like that of many vertebrates, is dominated by PRDM9-mediated hotspots, with artificial intelligence-based models predicting an intricate 3D-structure of the species-specific PRDM9-DNA binding complex *in silico*. Apart from providing novel insights into the population genetics of common marmosets, given the importance of the availability of fine-scale maps of mutation and recombination for evolutionary inference, this work will also serve as a valuable resource to aid future genomic research in this widely studied system.

## INTRODUCTION

The introduction of new genetic variation through the process of mutation and its reorganization through crossover and non-crossover events — the two possible outcomes of meiotic recombination — are key evolutionary mechanisms that influence genomic diversity. Both processes are known to be highly variable across species, with rates differing between populations, individuals, and sites within a genome (see the reviews by Baer et al. 2007; Lynch 2010; Hodgkinson and Eyre-Walker 2011; Pfeifer 2020b for an overview of mutation rate variation and a discussion of its genetic determinants, and the reviews by Ritz et al. 2017; Stapley et al. 2017; Johnston 2024 for an overview of recombination rate variation). Importantly, the uneven distribution of mutation and recombination rates across genomes can profoundly influence interactions amongst other evolutionary processes; for instance, heterogeneity in these underlying rates may substantially modify the effects of selection at linked sites, and thereby modify expectations of both levels and patterns of genetic variation (reviewed by Charlesworth and Jensen 2021, 2022). Additionally, relying on simplified, species-averaged rates of mutation and recombination — as is common practice in many evolutionary inference applications — has been shown to potentially lead to mis-inference in downstream analyses, including those estimating population history and the distribution of fitness effects (Dapper and Payseur 2018; Samuk and Noor 2022; Ghafoor et al. 2023; Soni et al. 2024; Soni and Jensen 2025). Yet, despite their crucial importance, both processes remain relatively poorly characterized in many vertebrates.

There are two primarily classes of approach for estimating both mutation and recombination rates in primates and other large organisms. Direct approaches utilize genomic data from parent-offspring trios or multi-generation pedigrees in order to detect *de novo* mutations (reviewed by Pfeifer 2020b, and see also Pfeifer 2021), or contemporary crossover and non-crossover events occurring between generations (reviewed by Clark et al. 2010). The resolution of these pedigree-based inference approaches at the genome scale remains relatively coarse due to the limited number of *de novo* mutations and meiotic exchanges that can be observed in the small number of generations characterizing the sample. Hence, given the hundreds to thousands of pedigreed individuals required to obtain high resolution maps, most studies typically yield only a genome-wide estimate over the limited generational span studied, rather than providing detailed information about the finer scale landscapes. Moreover, due to the extensive sample and sequencing requirements, direct approaches tend to be labor-intensive and costly, limiting their application, particularly in organisms for which resources remain limited. Indirect approaches, on the other hand, rely on population genetic theory and information about the genealogy of the sample over longer evolutionary timescales. Indirect estimates of mutation rate, for example, rely on species-level divergence data, based on the observation that the neutral mutation rate is equal to the neutral divergence rate (Kimura 1968). This allows for the inference of historically averaged mutation rates from phylogenetic sequence data in neutrally-evolving genomic regions. Although in principle straightforward, this approach is limited by the availability of high-quality genome annotations necessary to identify neutrally-evolving regions — which are lacking for many non-model organisms — and is often accompanied by significant uncertainties related to divergence and generation times, typically resulting in a range of possible mutation rates (see e.g., Soni et al. 2025c). Indirect recombination rate inference relies on polymorphism rather than divergence data, analyzing unrelated individuals in order to estimate historical sex-averaged rates of recombination based on the extent of observed linkage disequilibrium (LD) in the genome (reviewed by Stumpf and McVean 2003; Peñalba and Wolf 2020). For this reason, these methods are sensitive to other evolutionary forces shaping LD (Dapper and Payseur 2018; Samuk and Noor 2022) and it is thus crucial to account for the underlying population history when performing such analyses (Johri et al. 2020, 2022; Jensen 2023). Yet, despite these caveats, these approaches are also uniquely capable of providing a fine-scale, genome-wide mapping of underlying sex-averaged rates over many generations necessary for evolutionary analyses (see the discussion in Johri et al. 2022).

Within primates specifically, high-quality, fine-scale rate estimation has generally been focused upon humans and their closest relatives, the great apes, as well as a handful of species of biomedical or conservation interest (e.g., Kong et al. 2002; Auton et al. 2012; Stevison et al. 2016; Pfeifer 2020a; Xue et al. 2020; Wall et al. 2022; Versoza et al. 2024, 2025a,b; Soni et al. 2025c; and see the discussion of Soni et al. 2025d). In the primates studied to date as well as in numerous other organisms, meiotic recombination has been found to be concentrated in hotspots, the location of which is primarily determined by the zinc-finger protein PRDM9 (Baudat et al. 2010; Myers et al. 2010; Parvanov et al. 2010). Notably, PRDM9 has evolved rapidly across primates, showing a high variability in the number of zinc-fingers as well as their nucleotide contact residues, resulting in differences in the predicted nucleotide binding sequence and, consequently, in the hotspot positioning even between closely-related species (reviewed by Stapley et al. 2017; Lorenz and Mpaulo 2022; Johnston 2024; and see Schwartz et al. 2014 for a characterization of the allelic diversity in PRDM9 zinc finger domains of different primate species). As many evolutionary and biomedical studies require knowledge of the fine-scale patterns of recombination across the genome — such as rare disease gene mapping and genome-wide association studies that rely on patterns of LD to detect associations between genetic variants and phenotypic traits — and given the extensive variation in the distribution of recombination observed, it is thus important to obtain detailed maps characterizing the recombination frequency specific to the organism of interest. Yet, deeper in the primate clade, high-quality estimates of fine-scale mutation and recombination rates, or hotspot mapping, remain sparse — particularly across New World monkeys (platyrrhines) and prosimians (strepsirrhines) — limiting both our understanding of primate evolution in general and the accuracy and resolution of clinically-relevant genomic studies in primate model systems specifically.

In order to extend this inference, we here examine the common (or white-tufted-ear) marmoset, *Callithrix jacchus*, a relatively abundant platyrrhine monkey of the Callitrichidae family. As one of the smallest statured anthropoids, common marmosets have risen in biomedical prominence over the past two decades particularly as a commonly used model for the study of aging, neurodegeneration, and neurodevelopment disorders, as well as autoimmune and infectious disease dynamics (see e.g., the reviews by Miller et al. 2016; Philippens and Langermans 2021; Han et al. 2022), owing largely to their early sexual maturity (∼15 to 18 months of age), short gestation period (∼145 days), and high fecundity (birthing up to four offspring in a single pregnancy, with pregnancies occurring biannually). Interestingly, a notable peculiarity of marmoset biology, and that of several other callitrichines, is the frequent ovulation of two (or more) ova per cycle (Tardif and Jaquish 1997), resulting in common dizygotic (fraternal) twin births (Ward et al. 2014), and the exchange of hematopoietic stem cells during embryonic development via anastomoses in a single shared placenta, causing sibling chimerism (Hill 1932; Wislocki 1939; Benirschke et al. 1962; Gengozian et al. 1969). As a result, samples of blood and many other tissues obtained from marmosets have been observed to contain a mixture of genetic material originating from both the sampled individual and their unsampled littermate (Ross et al. 2007; Sweeney et al. 2012; del Rosario et al. 2024; and see the commentary by Chiou and Snyder-Mackler 2024). Notably, recent genomic analyses have demonstrated that the high frequency of twinning together with chimeric sampling shape both levels and patterns of observed genetic variation — including LD — in this species, with Soni et al. (2025b) finding that a neglect of these biological factors can result in significant mis-inference of population history. Yet, the impact of these unusual reproductive dynamics on other types of population genomic inference remains unexplored to date.

Although a direct mutation rate point estimate exists for common marmosets (0.43 × 10^-8^ per base pair per generation; Yang et al. 2021), this estimate was based on a single trio and, although the sampled tissues (muscle, liver, and spleen) are expected to exhibit lower levels of chimerism than blood, the authors did not correct for the effects of chimerism. In this instance, the potential ‘counting’ of twin-shared variants in a single individual sampled from a triplet may be expected to confound mutation counts and thus calculated rates. To lower the potential impact of chimerism on their estimate, the authors applied highly stringent filter criteria; however, this in turn may be expected to impact the overall number of genuine *de novo* mutations identifiable in their study. More recently, Mao et al. (2024) inferred a much-increased human-marmoset neutral divergence relative to the closely-related owl monkey-human divergence. In this case, the long-term neutral divergence rate would not be expected to be impacted by twinning and chimeric sampling (i.e., neutral divergence will be driven by the neutral mutation rate, *μ*). In contrast to these studies of mutation, to the best of our knowledge, no fine-scale genetic map yet exists for the species. Helpfully, utilizing novel population genomic data from individuals sequenced to high-coverage and annotated at the gene-level — and directly accounting for twinning and chimerism — Soni et al. (2025b) recently described a well-fitting population history consisting of a rapid reduction in population size roughly 7,000 years ago, followed by a modest recovery — a necessary prerequisite to interpret patterns of genetic variation and LD. Here, we utilize this modelling framework accounting both for this unique reproductive biology as well as population history, combined with high-quality whole-genome sequencing data, in order to quantify the rates and characterize the landscape of mutation and recombination in the common marmoset. Beyond providing valuable biological insights into the location and frequency of mutation and recombination in this New World monkey, these fine-scale maps will play a crucial role in enhancing future biomedical and evolutionary analyses integrating the significant nuance of accurate patterns of underlying genomic rate heterogeneity, thus improving our ability to study heritable genetic disorders in this widely used model system. In this effort, we have also uniquely characterized the effects of twinning and chimeric sampling on indirect rate estimation more generally and describe the underlying impact on patterns of LD — insights which will be beneficial to future research focused on organisms characterized by chimerism.

## MATERIALS AND METHODS

### Animal subjects

Animals were maintained in accordance with the guidelines of the Harvard Medical School Standing Committee on Animals and the Guide for Care and Use of Laboratory Animals of the Institute of Laboratory Animal Resources, National Research Council. All samples were collected during routine veterinary care under approved protocols.

### Whole genome sequencing

Our study was based on genomes of 15 captive common marmosets (*C. jacchus*) from the colony previously housed at the New England Primate Research Center. Whole genome sequencing data (150 bp paired-end reads) was generated on a DNBseq platform at the Beijing Genomics Institute (BGI Group, Shenzhen, China), targeting a genome-wide average coverage of ∼35X. Genetic analysis confirmed that the selected individuals were largely unrelated, sharing no more than 1/16 of their DNA with any other individual from the colony included in this study.

### Read mapping, variant calling, and filtering

Following best practices in the field (Pfeifer 2017), we trimmed adapters, polyX tails, and low-quality ends from the raw reads using SOAPnuke v.1.5.6 with BGI-recommended parameter settings (i.e., ’ *-n* 0.01 *-l* 20 *-q* 0.3 *-A* 0.25 *--cutAdaptor -Q* 2 *-G --polyX --minLen* 150’; Chen et al. 2018) before mapping them to the species-specific reference genome (mCalJa1.2.pat.X; GenBank accession number: GCA_011100555.2; Yang et al. 2021) using BWA-MEM v.0.7.17 (Li 2013). To avoid potential biases during variant calling and genotyping, we marked duplicates in the mapped reads using the Genome Analysis Toolkit (GATK) v.4.2.6.1 *MarkDuplicates* (van der Auwera and O’Connor 2020). From these high-quality, duplicate-marked reads (’ *--minimum-mapping-quality* 40 ’), we called variants for each individual using the GATK *HaplotypeCaller* in the ’ *--pcr-indel-model* NONE ’ mode as the data was obtained from PCR-free libraries. We then combined individual-level variant call sets and jointly genotyped them using GATK’s *CombineGVCFs* and *GenotypeGVCFs*, respectively. In order to obtain information regarding the genomic regions accessible to the study, we emitted reference confidence scores at each locus (’ *--emit-ref-confidence* BP_RESOLUTION ’) and included genotypes of both variant and invariant loci in the output (’ *--include-non-variant-sites* ’). Lastly, we separated the jointly-genotyped call set into biallelic single nucleotide polymorphisms (SNPs) and invariant loci with genotype information available in all individuals (’ *AN* = 30 ’), with downstream analyses focusing on the autosomes (i.e., chromosomes 1-22) only.

Due to the lack of a well-curated, experimentally-validated dataset that could be harnessed for filtering variants via a machine-learning framework, we used the GATK *VariantFiltration* tool to quality-control the discovered loci following the developers’ hard filtering recommendations (i.e., *QD* < 2.0, *QUAL* < 30.0, *MQ* < 40.0, *FS* > 60.0, *SOR* > 3.0, *MQRankSum* < -12.5, *ReadPosRankSum* < -8.0). As spurious variation can complicate the estimation of recombination rates from patterns of LD observed in population genomic data, we applied several additional stringent filtering criteria to limit the number of false positives in our data set, following the methodologies established in prior research (e.g., Auton et al. 2012; Stevison et al. 2016; Pfeifer 2020a; Soni et al. 2025c). In brief, as a skewed read coverage often results in poorly supported genotypes, we first used GATK’s *SelectVariants* to remove loci located within genomic regions exhibiting less than half, or more than twice, the individual’s genome-wide average coverage. Second, we excluded variants that were tightly clustered (using GATK’s *VariantFiltration* with ’ *--cluster-size* 3 ’ and ’ *--cluster-window-size* 10 ’) as well as those exhibiting high levels of heterozygosity (using VCFtools v.0.1.14 to filter out variants that have a Hardy-Weinberg equilibrium exact test *P*-value < 0.01; Danecek et al. 2011), a common indicator of local genome mis-assembly. Lastly, we used BEDTools v.2.30.0 (Quinlan et al. 2010) to eliminate regions blacklisted by the ENCODE Project Consortium due to their poor performance in next-generation sequencing (wgEncodeDacMapabilityConsensusExcludable.bed and wgEncodeDukeMapabilityRegionsExcludable.bed; Amemiya et al. 2019) by converting genomic coordinates between the human (hg38) and the common marmoset genome assemblies using the UCSC liftOver tool (Raney et al. 2024).

After filtering, the population-level dataset contained 7,198,428 autosomal biallelic SNPs in the accessible genome (Supplementary Table 1).

### Phasing

We estimated haplotypes from the population-level genotype data using BEAGLE v.5.5 (Browning et al. 2021), a progressive phasing algorithm that has been shown to be highly accurate (Williams et al. 2012). During phasing, a total of 16,701 loci (0.23%) became non-polymorphic due to a change in genotype and were thus excluded from further analyses.

### Inferring fine-scale rates of neutral divergence and mutation

In order to infer fine-scale rates of neutral divergence and mutation, we first used Cactus v.2.9.2 (Armstrong et al. 2020) to extract the 239 primate genomes (Kuderna et al. 2024) from the 447-way multiple species alignment (Zoonomia Consortium 2020) and update the common marmoset genome with the most recent reference genome for the species utilized in this study. To this end, we removed the common marmoset genome included in the multiple species alignment using the *halRemoveGenome* function and then extracted the alignment block consisting of the ancestral PrimateAnc232 genome and the genome of the closely related Wied’s black-tufted-ear marmoset (*C. kuhlii*) using the *hal2fasta* function. We then re-aligned the current reference genome for the common marmoset with the genomes of PrimateAnc232 and *C. kuhlii* using the previously inferred branch lengths and updated the alignment using the *halReplaceGenome* function. With this updated multiple species alignment on hand, we identified fixed single nucleotide differences along the marmoset lineage (i.e., between the common marmoset and the ancestral primate PrimateAnc232 genomes) using the *halSummarizeMutations* function. In contrast to the common marmoset genome, the genome of Wied’s marmoset is highly fragmented (293,512 scaffolds with a scaffold N50 of 15.1 kb totalling 2.6 Gb compared to 1,233 scaffolds with a scaffold N50 of 137 Mb totalling 2.9 Gb), thus we removed alignments less than 10 kb in length in order to avoid spurious and/or incomplete alignments that might artificially inflate estimates of neutral divergence and mutation. Additionally, in order to obtain fixed differences along the human-marmoset branch, we obtained the sub-alignment consisting of the common marmoset, human, and PrimatesAnc003 genomes from the multi-species alignment using the *cactus-hal2maf* function, converted this alignment back into HAL format (Hickey et al. 2013) using the *maf2hal* function, and then retrieved point mutations along the branch from *C. jacchus* to the ancestral PrimatesAnc003 using the *halBranchMutations* function. In order to obtain neutral substitutions between *C. jacchus* and *C. kuhlii* as well as between *C. jacchus* and humans, we excluded both variants known to segregate in any of the species (based on published population-level polymorphism data available for humans of Yoruban ancestry [1000 Genomes Project Consortium 2015] and common marmosets [Soni et al. 2025b]; note that no such population-level polymorphism data was available for Wied’s marmoset) as well as sites within 10 kb of functional regions (based on the protein-coding genes annotated in the common marmoset genome). Using these datasets, we then calculated neutral divergence at both the broad (genome-wide) scale and the fine scale (using 1 kb, 10 kb, 100 kb, and 1 Mb sliding windows with a step size of 500 bp, 5 kb, 50 kb, and 500 kb, respectively) by dividing the number of substitutions by the number of sites accessible to our study. To derive mutation rates, we divided the rates of neutral divergence between *C. jacchus* and *C. kuhlii* by the divergence time in generations. For this, we used three possible divergence time estimates — 0.59 million years ago (mya), 0.82 mya and 1.09 mya (based on the range of divergence times between *C. jacchus* and *C. kuhlii* inferred by Malukiewicz et al. 2021) — and two possible generation time estimates — 1.5 years and 2.0 years (Tardif et al. 2003; Okano et al. 2012; Schultz-Darken et al. 2016; Han et al. 2022).

### Inferring fine-scale rates of recombination

We used two different approaches to infer fine-scale rates of recombination: the demography-unaware estimator LDhat (McVean et al. 2002, 2004; Auton and McVean 2007) and its successor, the demography-aware estimator pyrho (Spence and Song 2019).

#### LDhat

We inferred genome-wide fine-scale recombination rates in the common marmoset using LDhat v.2.2 (McVean et al. 2002, 2004; Auton and McVean 2007). To this end, we first generated a lookup table containing the coalescent likelihoods for every two-locus haplotype configuration possible in our sample of 15 diploid individuals (i.e., 30 haploids: ’ *-n* 30 ’), using LDhat *complete* with the suggested maximum population-scaled recombination rate ρ of 100 (’ *-rhomax* 100 ’) and a grid size of 201 (’ *-n_pts* 201 ’) to improve accuracy. Next, we obtained region-based population-scaled estimates by running the *interval* function of LDhat with a block penalty of 5 (’ *-bpen* 5 ’) for 60 million iterations (’ *-its* 60000000 ’) using a sampling scheme of 40,000 iterations (’ *-samp* 40000 ’) across windows of 4,000 SNPs with a 200 SNP step size. After discarding the burn-in of the Monte Carlo Markov Chain using LDhat’s *stat* function (’ *-burn* 500’), we then combined the region-based estimates at the midpoint of each overlapping window to obtain estimates at the chromosome-scale. Following standard practices in the field (e.g., Auton et al. 2012; Pfeifer 2020a), we excluded large localized peaks in recombination rate (with ρ > 100 between adjacent SNPs) to minimize the impact of genome assembly errors leading to artificial breaks in LD. In total, we identified 1,352 such regions and masked them together with the 50 SNPs adjacent on each side by setting the recombination rate to 0 (masking a total of 42,348 SNPs, or 0.59%). Finally, we used *N_e_* based on the mean Θ observed in the empirical data to convert the population-scaled recombination rate to a per-generation recombination rate, assuming a per-site per-generation mutation rate of 0.81 ×10^-8^ as per Soni et al. 2025b.

#### pyrho

We also inferred genome-wide recombination rates using pyrho v.0.1.7 (Spence and Song 2019). In brief, as recommended by the developers, we first used pyrho’s *make_table* function to compute an approximate likelihood lookup table (’ *--approx* ’) for a 50% larger sample size (’ *-N* 45 ’) under the demographic model previously inferred by Soni et al. 2025b to account for historical population size changes in the species (i.e., a population size reduction approximately 3,500 generations ago, followed by an exponential population recovery) and then down-sampled this table to 15 diploid individuals (’ *-n* 30 ’) to match our empirical sample size. Next, we used the *hyperparam* function to determine suitable hyperparameter settings for pyrho and then ran the *optimize* function with the recommended window size (’ *--windowsize* 30 ’) and smoothness penalty (’ *--blockpenalty* 50 ’) to estimate genome-wide recombination rates. In all steps, we assumed a per-site per-generation mutation rate of 0.81 ×10^-8^ (’ *--mu* 0.81e-8 ’) as per Soni et al. 2025b to internally convert the population-scaled recombination rate to a per-generation recombination rate.

To account for the impact of twinning and chimerism (see "Evaluating the impact of twinning and chimerism on recombination rate inference" below), we rescaled the LDhat and pyrho estimates by multiplying the rates by a factor of 0.574 and 0.761, respectively.

### Evaluating the impact of twinning and chimerism on recombination rate inference

Both LDhat and pyrho rely on coalescent theory and the theoretical foundation provided by the Wright-Fisher model to infer fine-scale recombination rates from population genomic data. To evaluate the impact of twinning and chimerism — two model violations inherent to the biology of marmosets — on the recombination rate inference with these two approaches, we used SLiM 4.0.1 (Haller and Messer 2023) to simulate a population of marmoset individuals using the modelling framework recently described in Soni et al. (2025b). Specifically, we simulated 10 replicates of a 1 Mb genomic region in a population of marmosets under the demographic model of the species recently inferred by Soni et al. (2025b) — consisting of an ancestral population of 61,198 individuals that collapsed to 17,931 individuals 3,513 generations ago before recovering via exponential growth to a current day size of 33,830 individuals — assuming a mutation rate of 0.81 x 10^-8^ per base pair per generation and a recombination rate of 1.0 × 10^-8^ per base pair per generation (as per the rates used in Soni et al. 2025b). From each replicate, we sampled 15 chimeric individuals and estimated recombination rates using LDhat and pyrho (as described in "Inferring fine-scale rates of recombination").

### Inferring recombination hotpots

To infer the positions of recombination hotspots in the common marmoset, we used the software package LDhot v.0.4 (Auton et al. 2014). In brief, we first ran the *ldhot* function to identify putative recombination hotspots in 3 kb windows with a step size of 1 kb and a background window of 100 kb centered around the hotspot, using 1,000 simulations as recommended by the developers. Afterward, we combined significant windows to merge adjacent candidates using the *ldhot_summary* function with significance thresholds of 0.001 and 0.01 for calling and merging hotspots, respectively. To limit the number of spurious hotspots, we implemented a three-step filtering approach by combining the recommendations from the Great Ape Recombination Project (Stevison et al. 2016) with those of Brazier and Glémin (2024), filtering out any hotspot candidates with a width longer than 10 kb, an intensity lower than 4 or higher than 200, or a rate less than five times the chromosome-wide average rate.

### *In silico* prediction of PRDM9 binding motifs

We used the ZOOPS model implemented in MEME v.5.5.7 (Bailey and Elkan 1994) to identify 10-15 bp motifs present not more than once in each of the 1,000 hotspots with the highest intensity (including the flanking 500 bp to ensure that the entire hotspot is captured) while accounting for genomic background using cold spot regions matched for sequence length and GC-content. With these putative motifs on hand, we used FIMO v.5.5.7 to scan the complete set of hotspot windows for any occurrences and compared the frequency of each motif in the hotspot regions with that observed in 25,000 randomly sampled background regions using MOODS v.1.9.4.1 (Korhonen et al. 2009). We assessed the statistical significance by performing a Fisher’s exact test in R v.4.2.2 (R Core Team 2022).

### *In silico* characterization of PRDM9 binding

We used AlphaFold3 to predict the sequence-specific binding between PRDM9 and the putative PRDM9 binding motifs. To this end, we first retrieved the nucleotide sequence of PRDM9 from the common marmoset genome and aligned it against the PRDM9 protein sequence annotated in the human telomere-to-telomere assembly (T2T-CHM13v2.0; Nurk et al. 2022) using GeneWise v.2.4.1 (Birney et al. 2004) to visually inspect the sequence for completeness. Next, we used the ExPASy web server (Duvaud et al. 2021) to translate the nucleotide sequence into an amino acid sequence. We then used this translated amino acid sequence as input for a protein-protein BLAST (Altschul et al. 1990) search against the NCBI non-redundant protein sequences database (Supplementary Figure 1) and visualized the resulting phylogenetic tree using EMBL’s interactive Tree of Life (Letunic and Bork 2021), noting a high similarity of the query to PRDM9 sequences previously annotated in haplorrhines (Supplementary Figure 2). Next, we used InterPro (Blum et al. 2025) to predict protein domains within this sequence, confirming the presence of a Krueppel-associated box (KRAB) domain, a SSX repressor domain (SSXRD), a PR/SET domain, and a C2H2-type zinc finger array (Supplementary Figure 3). Finally, we ran AlphaFold3 (Abramson et al. 2024) by providing both the marmoset PRDM9 amino acid sequence and the putative PRDM9 binding motifs as input.

### Assessing the correlation of fine-scale rates of recombination with genomic features

To assess the correlation of fine-scale rates of recombination with different genomic features, we calculated a number of summary statistics across 100 kb windows along the 22 autosomes of the common marmoset genome — including nucleotide diversity (based on our marmoset population genomic data), divergence (based on the updated 447-way mammalian multiple species alignment as described in "Inferring fine-scale rates of neutral divergence and mutation"), GC-content and exon-content (both based on the annotations of the common marmoset assembly; Yang et al. 2021) — and calculated partial Kendall’s rank correlations across windows in which at least 50% of sites were accessible using R v.4.2.2 (R Core Team 2022).

## RESULTS AND DISCUSSION

### Population genomic data

Based on the genomes of 15 captive common marmoset (*C. jacchus*) individuals (seven females and eight males) sequenced to an average 35-fold coverage, we inferred fine-scale rates of neutral divergence, mutation, and recombination. In brief, using a mapping-based approach, we first identified 7.2 million SNPs across the autosomal genome with a transition-transversion ratio of 2.2 (Supplementary Table 1). To facilitate rate inference, we then estimated haplotypes from this population-level genotype data analogously to the 1000 Genomes (1000 Genomes Project Consortium 2015), PanMap (Auton et al. 2012), and Great Ape Recombination Maps (Stevison et al. 2016) projects, which previously generated fine-scale genetic maps for humans (*Homo sapiens*), Western and Nigerian chimpanzees (*Pan troglodytes verus* and *P. t. ellioti*), bonobos (*P. paniscus*), and Western gorillas (*Gorilla gorilla gorilla*).

### The landscape of neutral divergence and mutation in the common marmoset genome

We first extracted the 239 primate genomes (Kuderna et al. 2024) from the 447-way multiple species alignment (Zoonomia Consortium 2020) and updated the common marmoset genome to the current reference assembly available for the species (Yang et al. 2021). With this updated whole-genome alignment on hand, we identified fixed single nucleotide differences along the marmoset lineage, i.e., between the genome of the common marmoset and the genome of the closely related Wied’s black-tufted-ear marmoset (*C. kuhlii*). In contrast to the common marmoset genome generated by the Vertebrates Genomes Project using a combination of short-read (Illumina) and long-read (PacBio) sequencing data and scaffolded using high-throughput chromosome conformation capture (Hi-C) and Bionano optical data, the genome of Wied’s marmoset remains highly fragmented, containing nearly 300,000 scaffolds. In order to avoid spurious and/or incomplete alignments that might artificially inflate estimates of neutral divergence and mutation, we thus removed any alignments shorter than 10 kb in length. In order to obtain neutral substitutions between *C. jacchus* and *C. kuhlii*, we additionally masked both regions within 10 kb of known conserved or functional elements — thus, avoiding purifying and background selection effecting our analyses (Charlesworth et al. 1993) — and variants known to segregate in the species. Using this dataset, we then calculated neutral divergence across accessible sites at both the broad (genome-wide) scale and the fine scale (for additional details, see "Materials and Methods").

At the 1Mb-scale, we observed a neutral divergence rate of 9.85 × 10^-4^ along the marmoset lineage relative to the reconstructed ancestor (see Supplementary Figure 4 for the distributions of neutral divergence across a range of window sizes). To calculate the neutral mutation rate, we then drew the point estimate, upper and lower bounds of common marmoset divergence times relative to *C. kuhlii* (0.59 mya, 0.82 mya, and 1.09 mya; Malukiewicz et al. 2021), and generation times of 1.5 and 2.0 years (Tardif et al. 2003; Okano et al. 2012; Schultz-Darken et al. 2016; Han et al. 2022). Depending on the underlying assumptions, the mean neutral mutation rate varied from 0.14 × 10^-8^ mutations per base pair per generation (under a divergence time of 1.09 mya and a generation time of 1.5 years) to 0.33 × 10^-8^ mutations per base pair per generation (under a divergence time of 0.59 mya and a generation time of 2.0 years; Table 1 and see Figure 1a for density plots of neutral mutation rate estimates across this range of possible divergence and generation times and Figure 1b for the inferred genome-wide neutral mutation rates). Notably, these indirectly inferred neutral mutation rates are lower than the direct estimate of 0.43 × 10^-8^ mutations per base pair per generation obtained by Yang et al. (2021) from a single trio. While indirect phylogenetic approaches are naturally unable to observe strongly deleterious / lethal mutations that are purged from the population, the higher pedigree-based estimate observed by Yang et al. (2021) might also partially be attributed to unaccounted for chimerism present in their genomic data (with non-germline tissues sampled from a single individual of a triplet).

**Figure 1.**
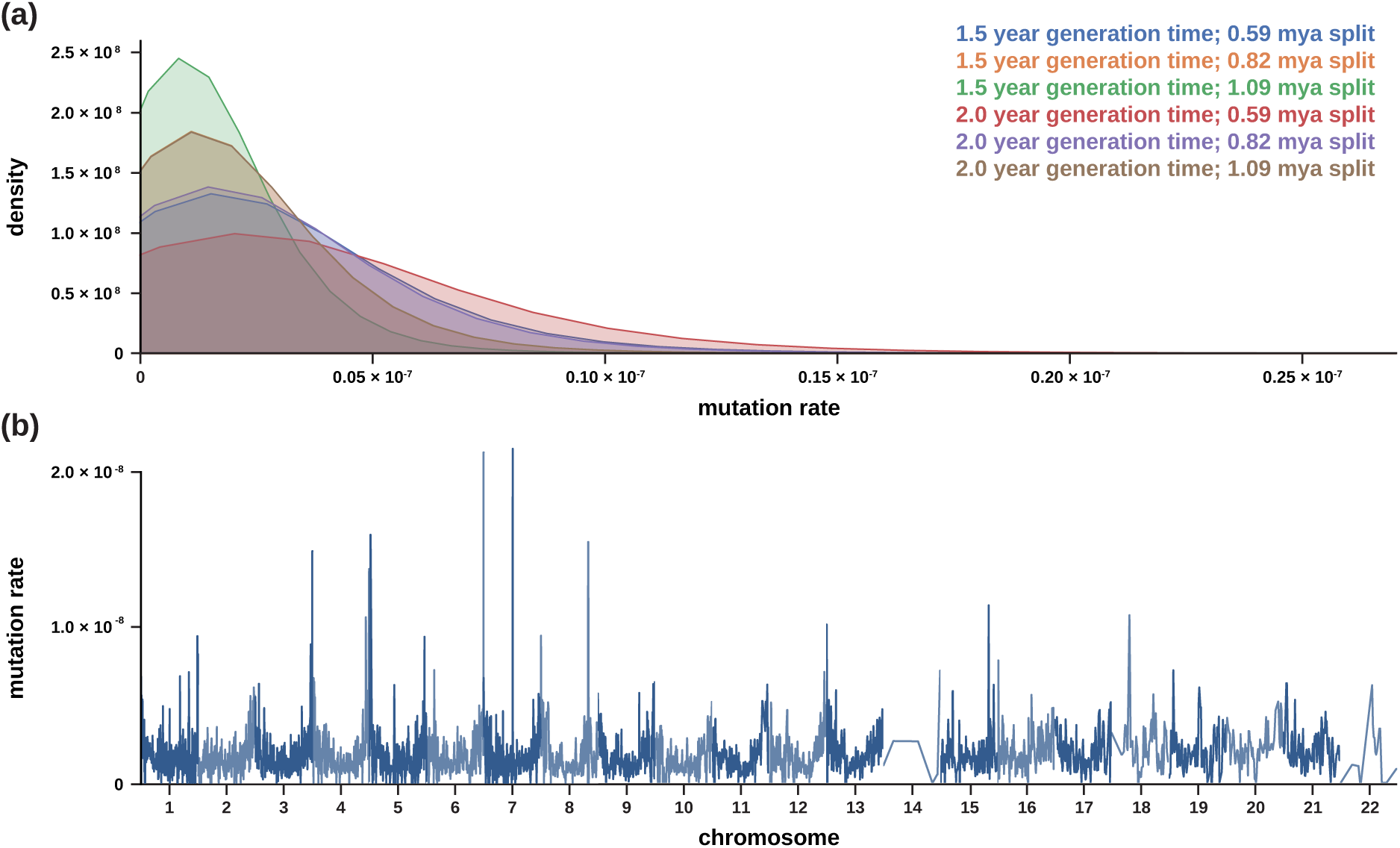
Fine-scale rates of neutral mutation. (a) Density plots of the per-site per-generation mutation rate implied by the neutral divergence for two possible generation times (1.5 years and 2.0 years; Tardif et al. 2003; Okano et al. 2012; Schultz-Darken et al. 2016; Han et al. 2022) and three possible divergence times between the common marmoset (*C. jacchus*) and the closely related Wied’s black-tufted-ear marmoset (*C. kuhlii*) (0.59 million years ago [mya], 0.82 mya, and 1.09 mya; Malukiewicz et al. 2021). (b) Genome-wide per-site per-generation neutral mutation rates for genomic windows of size 1 Mb, with a 500 kb step size (and see Supplementary Figure 5 for the heterogeneity in neutral mutation rates across all autosomes). Neutral mutation rates were estimated from the rates of neutral divergence observed in >10kb-long alignments between *C. jacchus* and *C. kuhlii* (note that due to the limited number of such alignments on chromosomes 14 and 22, neutral mutation rate estimates for these two autosomes are relatively coarse).

**Table 1.**
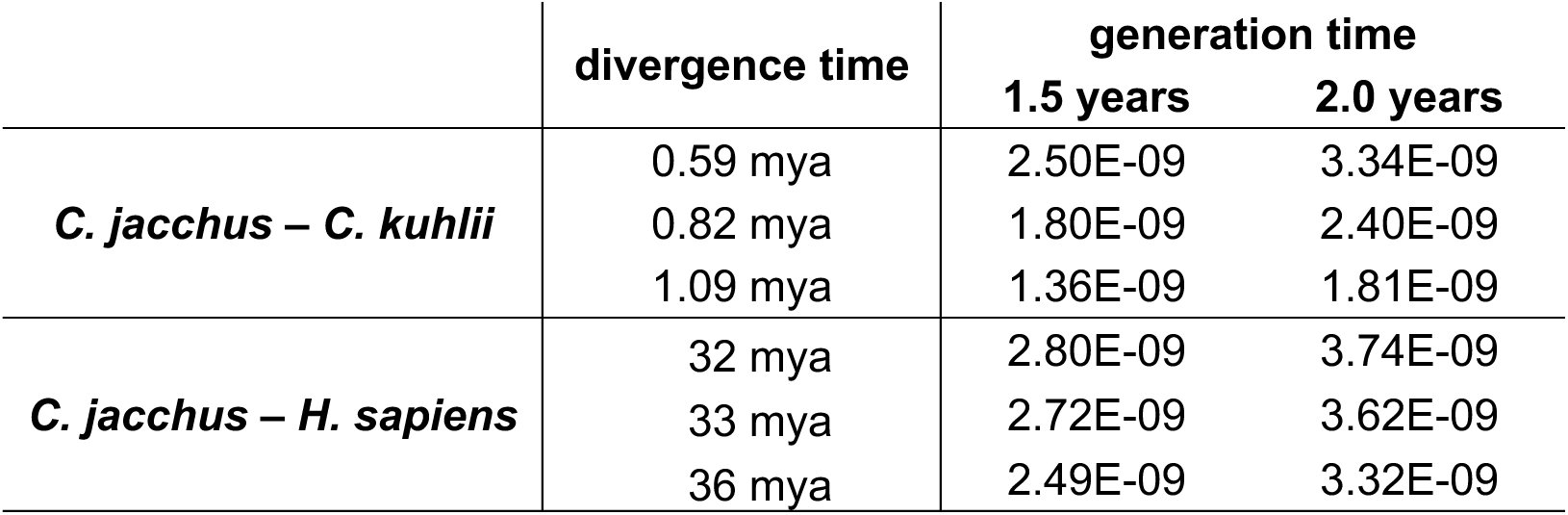
Inferred rates of neutral mutation. Comparison of indirectly inferred mean neutral mutation rate estimates based on *C. jacchus*–*C. kuhlii* divergence (mean neutral divergence rate: 0.0029) and *C. jacchus*–*H. sapiens* divergence (0.06) for three possible divergence times (0.59 million years ago [mya], 0.82 mya, and 1.09 mya for *C. jacchus*–*C. kuhlii* [Malukiewicz et al. 2021]; 32 mya, 33 mya, and 36 mya for *C. jacchus*–*H. sapiens* [Glazko and Nei 2003]) and two possible generation times (1.5 years and 2.0 years; Tardif et al. 2003; Okano et al. 2012; Schultz-Darken et al. 2016; Han et al. 2022).

Given that the divergence time between *C. jacchus* and *C. kuhlii* is relatively short, we also generated fine-scale divergence estimates along the human-marmoset branch to obtain information over longer evolutionary time scales. Notably, comparisons between the mutation rate estimates indirectly obtained from the *C. jacchus*–*C. kuhlii* divergence and the neutral divergence estimates based on *C. jacchus*–*H. sapiens* alignments demonstrated a significant positive correlation at the fine scale (*r* = 0.235, *P*-value = 1.71E-59; and see Supplementary Figure 5 for the heterogeneity in neutral divergence and mutation rates across each autosome), providing additional confidence in our estimates. Moreover, mutation rates inferred using the divergence along the human-marmoset branch are highly similar to those obtained using *C. jacchus*–*C. kuhlii* divergence, ranging from 0.25 × 10^-8^ mutations per base pair per generation (under a divergence time of 36 mya and a generation time of 1.5 years) to 0.37 × 10^-8^ mutations per base pair per generation (under a divergence time of 32 mya and a generation time of 2.0 years; Table 1). Although it is unlikely that the generation time of common marmosets has remained constant over the >30 million years separating the two species (Glazko and Nei 2003), it is nevertheless encouraging that the marmoset-based and human-marmoset-based estimates are in such close correspondence.

Taking the opposite approach, we inferred marmoset divergence times based on the mean neutral divergence rate observed in the empirical data and the previously published pedigree-based mutation rate estimates for both common marmosets (0.43 × 10^-8^ mutations per base pair per generation; Yang et al. 2021) as well as the closely related (but non-chimeric) owl monkeys (0.81 × 10^-8^ mutations per base pair per generation; Thomas et al. 2018) for which a larger number of trios were available. The estimated divergence times ranged from 0.18 mya (under a per-site per-generation mutation rate of 0.81 × 10^-8^ and a generation time of 1.5 years) to 0.49 mya (under a per-site per-generation mutation rate of 0.43 × 10^-8^ and a generation time of 2 years) (Supplementary Table 2). Notably, these whole-genome divergence times based on the marmoset and owl monkey mutation rates are considerably lower than those inferred by Malukiewicz et al. (2021) from mitochondrial DNA, due to the mutation rates being considerably higher than those inferred in this study.

### The landscape of recombination in the common marmoset genome

We used two different approaches to infer fine-scale rates of recombination — the demography-unaware estimator LDhat (McVean et al. 2002, 2004; Auton and McVean 2007) and its successor, the demography-aware estimator pyrho (Spence and Song 2019) — both of which rely on coalescent theory and the theoretical foundation provided by the Wright-Fisher (WF) model to infer fine-scale recombination rates from population genomic data.

Unlike most primates, twinning and chimerism are the norm rather than the exception in marmosets (Hill 1932; Wislocki 1939; Benirschke et al. 1962; Ward et al. 2014). To evaluate the impact of these two violations of the WF model inherent to the biology of marmosets, we first assessed the performance of the two recombination rate estimators on simulated data. Using the framework recently described in Soni et al. (2025b), we modelled twinning and chimerism from hematopoietic stem cells (as observed in blood samples) by first simulating a non-WF model in SLiM (Haller and Messer 2023) in which pairs of marmoset individuals reproduce each generation to give birth to non-identical twins and then combining their genotypes post-simulation to mimic a single chimeric individual (for additional details, see "Materials and Methods"). More specifically, we simulated genomic regions of 1 Mb under the demographic model of the population inferred by Soni et al. (2025b) — consisting of an ancestral population of ∼60k individuals that experienced a population decline before recovering to about half of its original size — using a non-WF model with twinning only as well as with both twinning and chimeric sampling, assuming a constant per-site per-generation mutation rate of 0.81 x 10^-8^ and recombination rate of 1 cM/Mb (as per the rates used in Soni et al. 2025b), and sampling 15 chimeric individuals to match our empirical data. To aid the interpretation of the inference results, we additionally performed simulations under a standard WF model (i.e., without twinning or chimerism) for comparison. We then inferred the genome-wide recombination rates in the simulated data using LDhat and pyrho. In contrast to pyrho, LDhat outputs the population-scaled recombination rate (ρ) and we thus used the effective population size (*N_e_*) based on the mean nucleotide diversity (Θ) observed in each simulation scenario to obtain a per-generation recombination rate.

Our simulations highlighted that both LDhat and pyrho tend to underestimate genome-wide recombination rates inferred under the marmoset demographic model (Figure 2a), likely due to the recent population contraction and subsequent exponential expansion having resulted in a ρ at the time of sampling that is substantially different from that during most of the population history. This observation is consistent with the recent findings of a simulation study conducted by Dutheil (2024) which highlighted that classical LD-based approaches tend to underestimate recombination rates in populations characterized by population size declines and recent growth, particularly when sample sizes are moderate as is the case here (see Figures 1 and 2 in Dutheil 2024; and see their Figure 4 for the increased mis-inference in the presence of gene conversion). In contrast, both approaches overestimate the per-generation recombination rate when twinning is common, as populations in which non-singleton births are the predominant mode of reproduction are characterized by both an increased LD and decreased *N_e_* relative to the standard expectations. Chimerism, on the other hand, has the opposite effect, with chimeric sampling expected to both break up haplotypes — as evidenced by the reduced mean *r^2^* (a measure of LD) — and increase haplotype diversity due to the generation of novel haplotypes relative to the standard expectations (Figure 2b).

**Figure 2:**
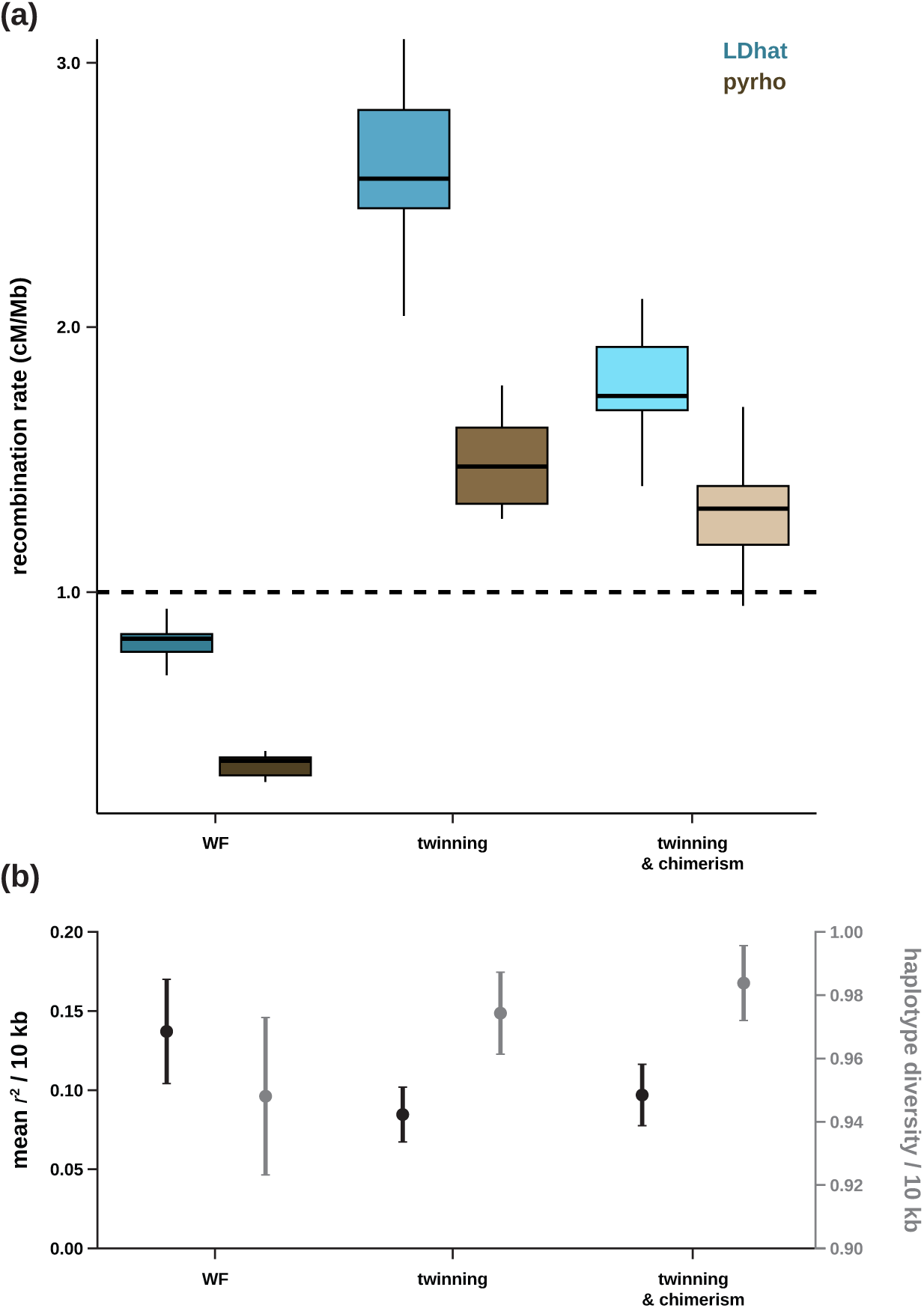
Performance of recombination rate estimators under twinning and chimerism. (a) Performance of the demography-unaware recombination rate estimator LDhat (shown in blues) and the demography-aware recombination rate estimator pyrho (shown in browns) under the demographic history inferred for the population by Soni et al. (2025b) using a Wright-Fisher (WF) model and a non-WF framework that models twinning and chimerism — two model violations inherent to the biology of marmosets. The dashed line depicts the recombination rate that was used in the simulations (1 cM/Mb). (b) Mean *r^2^* (shown in black; left y-axis) and haplotype diversity (gray; right y-axis) calculated over 10 kb windows (with a 5 kb step size) across 10 simulation replicates under the demographic history inferred for the population by Soni et al. (2025b) using a WF model and a non-WF framework that models twinning and chimerism. Data points represent the mean value, whilst confidence intervals represent the standard deviation.

Re-scaling the empirically observed recombination rate estimates to take into account the mis-inference caused by twinning and chimerism yielded an average sex-averaged genome-wide recombination rate of 0.91 ± 0.83 cM/Mb (100 kb windows) in common marmosets (Figure 3a), with sex-averaged crossover rates ranging from 0.65 cM/Mb for one of the longest autosomes (chromosome 4) to 1.29 cM/Mb for one of the shortest autosomes (chromosome 22) (Supplementary Figure 6). Although the rates inferred here for common marmosets are in the same range as those previously reported in the great apes (e.g., 1.32 ± 1.40 cM/Mb in humans [International HapMap Consortium 2007], with an average rate of 0.945 cM/Mb in males and 1.518 cM/Mb in females [Halldorsson et al. 2019], and ∼1.19 cM/Mb in chimpanzees, bonobos and gorillas [Stevison et al. 2016]), rates previously estimated for biomedically-relevant Old World monkeys are substantially lower (e.g., 0.43 ± 0.33 cM/Mb in rhesus macaques [Xue et al. 2016] and 0.43 ± 0.44 cM/Mb in vervet monkeys [Pfeifer 2020a]) — however, it should be noted that many of the earlier studies did not account for the potentially confounding effects of population demography during inference (see discussions of Dapper and Payseur 2018; Johri et al. 2022), thus hindering the interpretation of the biological differences in observed rates.

**Figure 3:**
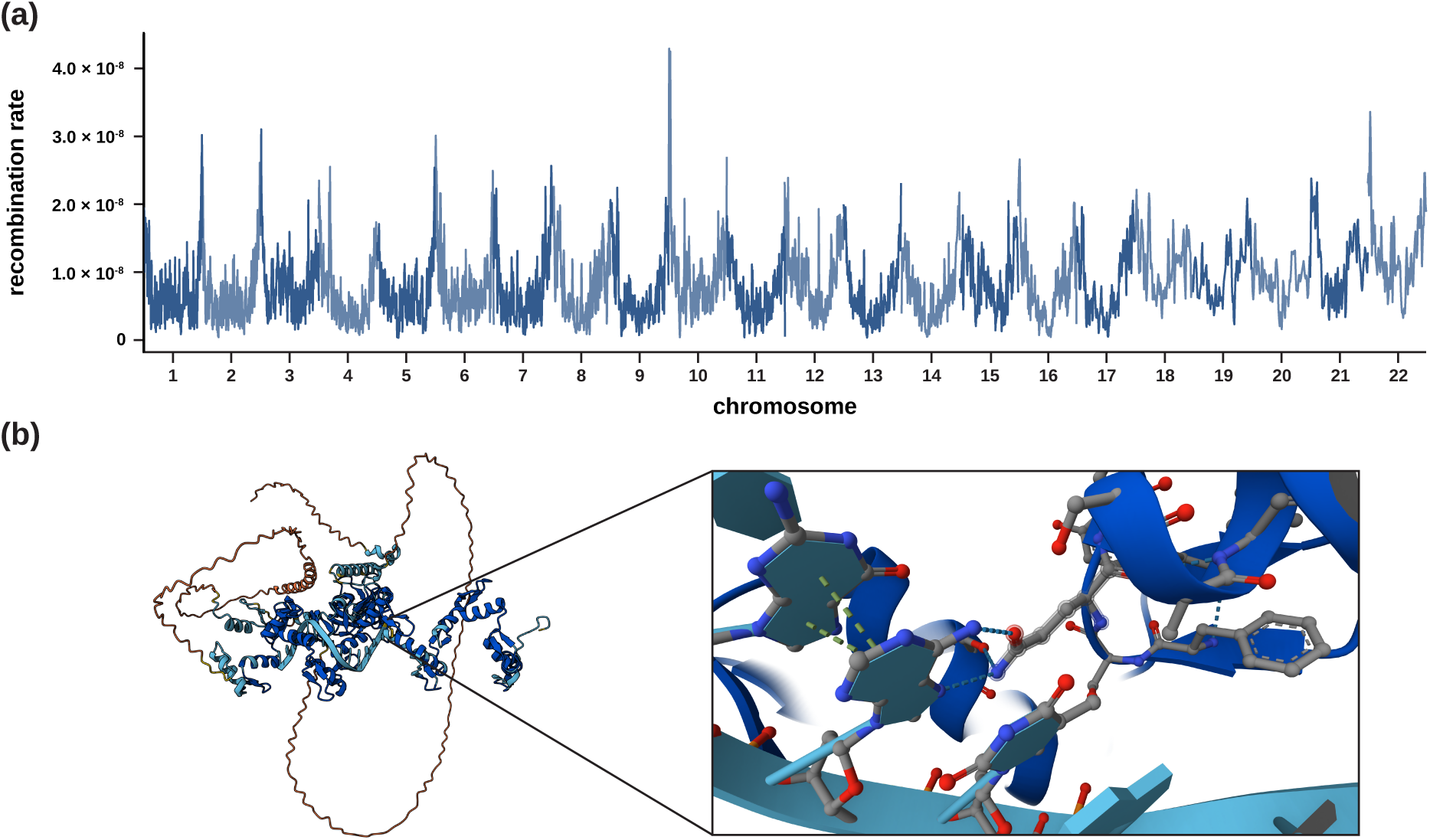
Fine-scale rates of recombination. (a) Genome-wide per-site per-generation recombination rates as inferred by LDhat for genomic windows of size 1Mb, with a 500kb step size (and see Supplementary Figure 7 for recombination rate heterogeneity across all autosomes and Supplementary Figure 10 for the genome-wide recombination rates inferred by pyrho). (b) *In silico* binding prediction between PRDM9 and the putative binding sites. Predictions were performed using AlphaFold3 (predictions are provided for non-commercial use only, under and subject to AlphaFold Server Output Terms of Use found under alphafoldserver.com/output-terms). Per-atom confidence scores are color-coded with very high confidence predicted structures shown in dark blue (pLDDT > 90), confidently predicted structures in turquoise (90 > pLDTT > 70), low confidence predicted structures in yellow (70 > pLDTT > 50), and very low confidence predicted structures in orange (pLDTT < 50).

In addition to the nearly two-fold variation between autosomes, the recombination landscape in common marmosets — like that of many other primates (e.g., Spencer et al. 2006; Auton et al. 2012; Pfeifer and Jensen 2016; Stevison et al. 2016; Pfeifer 2020a; Wall et al. 2022; Versoza et al. 2024; Soni et al. 2025c) — is highly heterogenous, with a pronounced enrichment in the distal (sub)telomeric regions and reduced rates within centromeric and pericentromeric regions (Figure 3a, and see Supplementary Figure 7 for recombination rate heterogeneity across all autosomes). Moreover, genome-wide recombination rates exhibit a strong positive association with nucleotide diversity (*r* = 0.49, *P*-value = 0) — highlighting the likely importance of selection at linked sites in shaping the landscape of genomic variation in the species (Soni et al. 2025a) — and, to a lesser extent, divergence (*r* = 0.06, *P*-value = 2.07E-17) and GC-content (*r* = 0.03, *P*-value = 6.90E-06), as may be expected if recombination in common marmosets exhibits mutagenic effects (Halldorsson et al. 2019; Hinch et al. 2023) and is prone to GC-biased gene conversion (reviewed by Duret and Galtier 2009) (Supplementary Figure 8). In concordance with PRDM9 directing recombination away from functional elements (Brick et al. 2012; Pratto et al. 2014), recombination rates significantly decrease within, and around the boundaries of, protein-coding genes (*r* = -0.03, *P*-value = 1.09E-04) compared to neighboring regions (Supplementary Figure 9). Notably, while these inferred patterns of recombination were highly similar between the two approaches (*r* = 0.57, *P*-value < 2.2E-16 at the 1kb-scale), the mean inferred rates of pyrho were much lower (0.25 ± 0.27 cM/Mb) as, despite accounting for historical fluctuations in population size, its internal conversion to the per-generation recombination rate does not account for the lower *N_e_* caused by twinning (Supplementary Figure 10).

### Characterization of recombination hotspots in the common marmoset genome

To gain insights into the landscape of recombination hotspots in the common marmoset genome, we inferred recombination hotspots using LDhot (Auton et al. 2014) — a statistical method implemented to directly process the recombination rate estimates obtained from LDhat. To obtain a high-quality dataset, we implemented a three-step filtering approach by combining the recommendations from the Great Ape Recombination Project (Stevison et al. 2016) with those of Brazier and Glémin (2024). First, as genuine recombination hotspots tend to be narrow (with ∼70-80% of recombination occurring in hotspots with a mean width of 2.3 kb in humans; 1000 Genomes Project Consortium 2010), we excluded hotspot candidates with a width larger than 10 kb. Second, as artificial breaks in LD resulting, for example, from mis-assembly and/or mis-alignment, can lead to localized peaks in recombination rate, we calculated the intensity of each hotspot candidate by dividing the recombination rate at its peak by the rate estimated for the 100 kb surrounding region and removed candidates with an intensity higher than 200. Additionally, due to the difficulty of distinguishing genuine low-intensity hotspots from background variation, we also applied a low-intensity filter, retaining only candidates with a minimum intensity of 4. Third, accounting for recombination rate variation across chromosomes to further narrow down the location of recombination hotspots, we split each remaining candidate region into windows of 2 kb with a step size of 1 kb, keeping only the hotspot windows exhibiting rates of at least five times the chromosome-wide average rate. This three-step filtering approach yielded 26,831 candidate hotspots — similar to the number of recombination hotspots initially identified in human populations (International HapMap Consortium 2005, 2007; Myers et al. 2005; 1000 Genomes Project Consortium 2010; Kong et al. 2012; though note the larger number of hotspots identified in subsequent, larger-scale studies, e.g., Halldorsson et al. 2019).

In order to identify sequences that predict potential PRDM9 binding, we searched the 1,000 hotspots with the highest intensity for consistent sequence motifs. To this end, we used the ZOOPS model implemented in MEME (Bailey and Elkan 1994) to identify degenerate motifs present not more than once in each hotspot region, while accounting for genomic background using cold spot regions matched for sequence length and GC-content. As sequence motifs underlying recombination hotspots in other primates tend to be short (e.g., 10-15 bp in humans, chimpanzees, and gorillas; Schwartz et al. 2014; Auton et al. 2012; Stevison et al. 2016), we limited the search to motifs with a minimum width of 10 bp and maximum width of 15 bp, keeping only motifs below a discovery threshold of 1e-05. Next, we used FIMO to scan the complete set of hotspot windows for occurrences of each of these motifs. As the predicted motifs are highly repetitive, we also performed permutation tests to compare the frequency of each motif in hotspot regions with that observed in 25,000 background regions randomly sampled from the common marmoset genome by matching the position weight matrix of each motif against the genomic sequences of each hotspot and cold spot region using the MOtif Occurrence Detection Suite (MOODS; Korhonen et al. 2009), requiring a *P*-value < 0.01 for a match. To further narrow down the list of candidates, we used AlphaFold3 (Abramson et al. 2024) to predict the sequence-specific binding between the marmoset PRDM9 sequence and the putative PRDM9 binding motifs. Interestingly, for the best candidate DNA binding motifs — the 15-mer GCTGGGATTACAGGC (*e*-value: 6.80E-54), present in 83.5% of the candidate hotspots and only 2.0% of the matching cold spots, and its degenerate generalization, the 15-mer GCTGGGAKYASWGGC (*e*-value: 5.40E-17) — AlphaFold3’s machine-learning guided model predicted a protein-DNA complex with high-confidence (ipTM = 0.94 and 0.96, respectively; with 1.0 being the maximum value of the ipTM performance metric), indicating a very well-defined protein-DNA interface and supporting PRDM9-DNA binding *in silico* (see Figure 3b for a visualization of the predicted binding complex). Notably, while this predicted degenerate PRDM9 binding motif shares several key features with motifs previously observed in other primates (including the C/G-rich terminal regions and its overall length), there are also species-specific differences (such as the A/T-rich internal region) as might be anticipated from the rapid evolution of the PRDM9 zinc finger array across the primate lineage.

## CONCLUSION

By investigating, and ultimately correcting for, the effects of twinning and sibling chimerism on the inference of fine-scale mutation and recombination rate maps, we have here described these landscapes in one of the most commonly used non-human primate models in research, *C. jacchus*. Using high-quality population genomic data, we found that the species exhibits relatively low neutral mutation rates, and rates of recombination within the range observed amongst other anthropoids. Moreover, like many vertebrates, the recombination landscape in common marmosets is dominated by PRDM9-mediated hotspots, and we have described a 3D-structure of the species-specific PRDM9-DNA binding complex *in silico*. Apart from providing novel insights into the population genetic processes shaping variation in common marmosets, these maps will also serve as a valuable resource for future studies in this biomedically important species — including in genome-wide association studies, polygenic risk score modelling, and genomic scans for targets of selection — with implications ranging from the improved study of neurodevelopment disorders to infectious disease dynamics.

## ACKNOWLEDGEMENTS

We would like to thank Eric J. Vallender for providing the marmoset samples used in this study, and the members of the Jensen and Pfeifer Labs for helpful comments and discussion. Library preparation and sequencing was performed at the Beijing Genomics Institute (BGI Group, Shenzhen, China). Computations were performed on the Sol supercomputer at Arizona State University (Jennewein et al. 2023).

## REFERENCES

1000 Genomes Project Consortium. 2010. A map of human genome variation from population-scale sequencing. Nature. 467(7319):1061–1073.

1000 Genomes Project Consortium. 2015. A global reference for human genetic variation. Nature. 526(7571):68–74.

Abramson J, Adler J, Dunger J, Evans R, Green T, Pritzel A, Ronneberger O, Willmore L, Ballard AJ, Bambrick J, et al. 2024. Accurate structure prediction of biomolecular interactions with AlphaFold 3. Nature. 630(8016):493–500.

Altschul SF, Gish W, Miller W, Myers EW, Lipman DJ. 1990. Basic local alignment search tool. J Mol Biol. 215(3):403–410.

Amemiya HM, Kundaje A, Boyle AP. 2019. The ENCODE blacklist: identification of problematic regions of the genome. Sci Rep. 9(1):9354.

Armstrong J, Hickey G, Diekhans M, Fiddes IT, Novak AM, Deran A, Fang Q, Xie D, Feng S, Stiller J, et al. 2020. Progressive Cactus is a multiple-genome aligner for the thousand-genome era. Nature. 587(7833):246–251.

Auton A, Fledel-Alon A, Pfeifer S, Venn O, Ségurel L, Street T, Leffler EM, Bowden R, Aneas I, Broxholme J, et al. 2012. A fine-scale chimpanzee genetic map from population sequencing. Science. 336(6078):193–198.

Auton A, McVean G. 2007. Recombination rate estimation in the presence of hotspots. Genome Res.17(8):1219–1227.

Auton A, Myers S, McVean G. 2014. Identifying recombination hotspots using population genetic data. arXiv [Preprint] 10.48550/arXiv.1403.4264

Baer CF, Miyamoto MM, Denver DR. 2007. Mutation rate variation in multicellular eukaryotes: causes and consequences. Nat Rev Genet. 8(8):619–631.

Bailey TL, Elkan C. 1994. Fitting a mixture model by expectation maximization to discover motifs in biopolymers. Proc Int Conf Intell Syst Mol Biol. 2:28–36.

Baudat F, Buard J, Grey C, Fledel-Alon A, Ober C, Przeworski M, Coop G, de Massy B. 2010. PRDM9 is a major determinant of meiotic recombination hotspots in humans and mice. Science. 327(5967):836–840.

Benirschke K, Anderson JM, Brownhill LE. 1962. Marrow chimerism in marmosets. Science. 138(3539):513–515.

Blum M, Andreeva A, Florentino LC, Chuguransky SR, Grego T, Hobbs E, Pinto BL, Orr A, Paysan-Lafosse T, Ponamareva I, et al. 2025. InterPro: the protein sequence classification resource in 2025. Nucleic Acids Res. 53(D1):D444–D456.

Birney E, Clamp M, Durbin R. 2004. GeneWise and Genomewise. Genome Res. 14(5):988–995.

Brazier T, Glémin S. 2024. Diversity in recombination hotspot characteristics and gene structure shape fine-scale recombination patterns in plant genomes. Mol Biol Evol. 41(9):msae183.

Brick K, Smagulova F, Khil P, Camerini-Otero RD, Petukhova GV. 2012. Genetic recombination is directed away from functional genomic elements in mice. Nature. 485(7400):642–645.

Browning BL, Tian X, Zhou Y, Browning SR. 2021. Fast two-stage phasing of large-scale sequence data. Am J Hum Genet. 108(10):1880–1890.

Charlesworth B, Jensen JD. 2021. Effects of selection at linked sites on patterns of genetic variability. Annu Rev Ecol Evol Syst. 52:177–197.

Charlesworth B, Jensen JD. 2022. How can we resolve Lewontin’s Paradox? Gen Biol Evol. 14(7):evac096.

Charlesworth B, Morgan MT, Charlesworth D. 1993. The effect of deleterious mutations on neutral molecular variation. Genetics. 134(4):1289–1303.

Chen Y, Chen Y, Shi C, Huang Z, Zhang Y, Li S, Li Y, Ye J, Yu C, Li Z, et al. 2018. SOAPnuke: a MapReduce acceleration-supported software for integrated quality control and preprocessing of high-throughput sequencing data. GigaScience. 7(1):1–6.

Chiou K, Snyder-Mackler N. 2024. Marmosets contain multitudes. eLife 13:e97866.

Clark AG, Wang X, Matise T. 2010. Contrasting methods of quantifying fine structure of human recombination. Annu Rev Genomics Hum Genet. 11:45–64.

Danecek P, Auton A, Abecasis G, Albers CA, Banks E, DePristo MA, Handsaker RE, Lunter G, Marth GT, Sherry ST, et al. 2011. The variant call format and VCFtools. Bioinformatics. 27(15):2156–2158.

Dapper AL, Payseur BA. 2018. Effects of demographic history on the detection of recombination hotspots from linkage disequilibrium. Mol Biol Evol. 35(2):335–353.

del Rosario RCH, Krienen FM, Zhang Q, Goldman M, Mello C, Lutservitz A, Ichihara K, Wysoker A, Nemesh J, Feng G, et al. 2024. Sibling chimerism among microglia in marmosets. eLife 13:RP93640.

Duret L, Galtier N. 2009. Biased gene conversion and the evolution of mammalian genomic landscapes. Annu Rev Genomics Hum Genet. 10:285–311.

Dutheil JY. 2024. On the estimation of genome-average recombination rates. Genetics. 227(2):iyae051.

Duvaud S, Gabella C, Lisacek F, Stockinger H, Ioannidis V, Durinx C. 2021. Expasy, the Swiss Bioinformatics Resource Portal, as designed by its users. Nucleic Acids Res. 49(W1):W216–W227.

Ghafoor S, Santos J, Versoza CJ, Jensen JD, Pfeifer SP. 2023. The impact of sample size and population history on observed mutational spectra: a case study in human and chimpanzee populations. Gen Biol Evol. 15(3):evad019.

Glazko GV, Nei M. 2003. Estimation of divergence times for major lineages of primate species. Mol Biol Evol. 20(3):424–434.

Gengozian N, Batson JS, Greene CT, Gosslee DG. 1969. Hemopoietic chimerism in imported and laboratory-bred marmosets. Transplantation. 8(5):633–652.

Halldorsson BV, Palsson G, Stefansson OA, Jonsson H, Hardarson MT, Eggertsson HP, Gunnarsson B, Oddsson A, Halldorsson GH, Zink F, et al. 2019. Characterizing mutagenic effects of recombination through a sequence-level genetic map. Science. 363(6425):eaau1043.

Haller BC, Messer PW. 2023. SLiM 4: multispecies eco-evolutionary modeling. Am Nat. 201:E127–E139.

Han HJ, Powers SJ, Gabrielson KL. 2022. The common marmoset – biomedical research animal model applications and common spontaneous diseases. Toxicol Pathol. 50(5):628–637.

Hickey G, Paten B, Earl D, Zerbino D, Haussler D. 2013. HAL: a hierarchical format for storing and analyzing multiple genome alignments. Bioinformatics. 29(10):1341–1342.

Hill JP. 1932. II. Croonian lecture - The developmental history of the primates. Phil Trans R Soc Lond B. 221(474–482):45–178.

Hinch R, Donnelly P, Hinch AG. 2023. Meiotic DNA breaks drive multifaceted mutagenesis in the human germ line. Science. 382(6674):eadh2531.

Hodgkinson A, Eyre-Walker A. 2011. Variation in the mutation rate across mammalian genomes. Nat Rev Genet. 12(11):756–766.

International HapMap Consortium. 2005. A haplotype map of the human genome. Nature. 437(7063):1299–1320.

International HapMap Consortium. 2007. A second generation human haplotype map of over 3.1 million SNPs. Nature. 449(7164):851–861.

Jennewein DM, Lee J, Kurtz C, Dizon W, Shaeffer I, Chapman A, Chiquete A, Burks J, Carlson A, Mason N, et al. 2023. The Sol supercomputer at Arizona State University. In Practice and Experience in Advanced Research Computing 2023: Computing for the Common Good (PEARC ’23). Association for Computing Machinery, New York, NY, USA, 296–301.

Jensen JD. 2023. Population genetic concerns related to the interpretation of empirical outliers and the neglect of common evolutionary processes. Heredity (Edinb). 130(3):109–110.

Johnston SE. 2024. Understanding the genetic basis of variation in meiotic recombination: past, present, and future. Mol Biol Evol. 41(7):msae112.

Johri P, Aquadro CF, Beaumont M, Charlesworth B, Excoffier L, Eyre-Walker A, Keightley PD, Lynch M, McVean G, Payseur BA, et al. 2022. Recommendations for improving statistical inference in population genomics. PLoS Biol. 20(5):e3001669.

Johri P, Charlesworth B, Jensen JD. 2020. Toward an evolutionarily appropriate null model: jointly inferring demography and purifying selection. Genetics. 215(1):173–192.

Kimura M. 1968. Evolutionary rate at the molecular level. Nature. 217(5129):624–626.

Kong A, Gudbjartsson DF, Sainz J, Jonsdottir GM, Gudjonsson SA, Richardsson B, Sigurdardottir S, Barnard J, Hallbeck B, Masson G, et al. 2002. A high-resolution recombination map of the human genome. Nat Genet. 31(3):241–247.

Korhonen J, Martinmäki P, Pizzi C, Rastas P, Ukkonen E. 2009. MOODS: fast search for position weight matrix matches in DNA sequences. Bioinformatics. 25(23):3181–3182.

Kuderna LFK, Ulirsch JC, Rashid S, Ameen M, Sundaram L, Hickey G, Cox AJ, Gao H, Kumar A, Aguet F, et al. 2024. Identification of constrained sequence elements across 239 primate genomes. Nature. 625(7996):735–742.

Letunic I, Bork P. 2021. Interactive Tree Of Life (iTOL) v5: an online tool for phylogenetic tree display and annotation. Nucleic Acids Res. 49(W1):W293–W296.

Li H. 2013. Aligning sequence reads, clone sequences and assembly contigs with BWA-MEM. arXiv [Preprint] DOI: 10.48550/arXiv.1303.3997

Lorenz A, Mpaulo SJ. 2022. Gene conversion: a non-Mendelian process integral to meiotic recombination. Heredity (Edinb). 129(1):56–63.

Lynch M. 2010. Evolution of the mutation rate. Trends Genet. 26(8):345–352.

Malukiewicz J, Cartwright RA, Curi NHA, Dergam JA, Igayara CS, Moreira SB, Molina CV, Nicola PA, Noll A, Passamani M, et al. 2021. Mitogenomic phylogeny of Callithrix with special focus on human transferred taxa. BMC Genomics. 22(1):239.

Mao Y, Harvey WT, Porubsky D, Munson KM, Hoekzema K, Lewis AP, Audano PA, Rozanski A, Yang X, Zhang S, et al. 2024. Structurally divergent and recurrently mutated regions of primate genomes. Cell. 187(6):1547–1562.e13.

McVean G, Awadalla P, Fearnhead P. 2002. A coalescent-based method for detecting and estimating recombination from gene sequences. Genetics. 160(3):1231–1241.

McVean GAT, Myers SR, Hunt S, Deloukas P, Bentley DR, Donnelly P. 2004. The fine-scale structure of recombination rate variation in the human genome. Science. 304(5670):581–584.

Miller CT, Freiwald WA, Leopold DA, Mitchell JF, Silva AC, Wang X. 2016. Marmosets: a neuroscientific model of human social behavior. Neuron. 90(2):219233.

Myers S, Bottolo L, Freeman C, McVean G, Donnelly P. 2005. A fine-scale map of recombination rates and hotspots across the human genome. Science. 310(5746):321–314.

Myers S, Bowden R, Tumian A, Bontrop RE, Freeman C, MacFie TS, McVean G, Donnelly P. 2010. Drive against hotspot motifs in primates implicates the PRDM9 gene in meiotic recombination. Science. 327(5967):876–879.

Nurk S, Koren S, Rhie A, Rautiainen M, Bzikadze AV, Mikheenko A, Vollger MR, Altemose N, Uralsky L, Gershman A, et al. 2022. The complete sequence of a human genome. Science. 376(6588):44–53.

Okano H, Hikishima K, Iriki A, Sasaki E. 2012. The common marmoset as a novel animal model system for biomedical and neuroscience research applications. Semin Fetal Neonatal Med. 17(6):336–340.

Parvanov ED, Petkov PM, Paigen K. 2010. PRDM9 controls activation of mammalian recombination hotspots. Science. 327(5967):835.

Peñalba JV, Wolf JBW. 2020. From molecules to populations: appreciating and estimating recombination rate variation. Nat Rev Genet. 21(8):476–492.

Pfeifer SP. 2017. From next-generation resequencing reads to a high-quality variant data set. Heredity (Edinb). 118(2):111–124.

Pfeifer SP. 2020a. A fine-scale genetic map for vervet monkeys. Mol Biol Evol. 37(7):1855–1865.

Pfeifer SP. 2020b. Spontaneous mutation rates. In Ho SYW (ed) The Molecular Evolutionary Clock. Theory and Practice. Springer Nature, pp. 35–44.

Pfeifer SP. 2021. Studying mutation rate evolution in primates-the effects of computational pipelines and parameter choices. GigaScience. 10(10):giab069.

Pfeifer SP, Jensen JD. 2016. The impact of linked selection in chimpanzees: a comparative study. Genome Biol Evol. 8(10):3202–3208.

Philippens IHCHM, Langermans JAM. 2021. Preclinical marmoset model for targeting chronic inflammation as a strategy to prevent Alzheimer’s disease. Vaccines (Basel). 9(4):388.

Pratto F, Brick K, Khil P, Smagulova F, Petukhova GV, Camerini-Otero RD. 2014. Recombination initiation maps of individual human genomes. Science. 346(6211):1256442.

Quinlan AR. 2014. BEDTools: the Swiss-army tool for genome feature analysis. Curr Protoc Bioinformatics. 47:11.12.1–34.

R Core Team. 2022. R: a language and environment for statistical computing. R Foundation for Statistical Computing, Vienna, Austria. https://www.R-project.org/

Raney BJ, Barber GP, Benet-Pagès A, Casper J, Clawson H, Cline MS, Diekhans M, Fischer C, Navarro Gonzalez J, Hickey G, et al. 2024. The UCSC Genome Browser database: 2024 update. Nucleic Acids Res. 52(D1):D1082–D1088.

Ritz RR, Noor MAF, Singh ND. 2017. Variation in recombination rate: adaptive or not? Trends Genet. 33(5):364–374.

Ross CN, French JA, Ortí G. 2007. Germ-line chimerism and paternal care in marmosets (*Callithrix kuhlii*). Proc Natl Acad Sci USA. 104(15):6278–6282.

Samuk K, Noor MAF. 2022. Gene flow biases population genetic inference of recombination rate. G3 (Bethesda). 12(11):jkac236.

Schultz-Darken N, Braun KM, Emborg ME. 2016. Neurobehavioral development of common marmoset monkeys. Dev Psychobiol. 58(2):141–158.

Schwartz JJ, Roach DJ, Thomas JH, Shendure J. 2014. Primate evolution of the recombination regulator PRDM9. Nat Commun. 5:4370.

Soni V, Jensen JD. 2025. Inferring demographic and selective histories from population genomic data using a two-step approach in species with coding-sparse genomes: an application to human data. G3 (Bethesda). 15:jkaf019.

Soni V, Pfeifer SP, Jensen JD. 2024. The effect of mutation and recombination rate heterogeneity on the inference of demography and the distribution of fitness effects. Genome Biol Evol. 16(2):evae004.

Soni V, Pfeifer SP, Jensen JD. 2025d. Recent insights into the evolutionary genomics of the critically endangered aye-aye (*Daubentonia madagascariensis*). Am J Primatol. 87(12):e70105.

Soni V, Versoza CJ, Pfeifer SP, Jensen JD. 2025a. Investigating the effects of chimerism on the inference of selection: quantifying genomic targets of purifying, positive, and balancing selection in common marmosets (*Callithrix jacchus*). Heredity (Edinb). 134(10-11):645–657.

Soni V, Versoza CJ, Terbot JW, Jensen JD, Pfeifer SP. 2025c. Inferring fine-scale mutation and recombination rate maps in aye-ayes (*Daubentonia madagascariensis*). Ecol. Evol. 15(11):e72314.

Soni V, Versoza CJ, Vallender EJ, Jensen JD, Pfeifer SP. 2025b. Accounting for chimerism in demographic inference: reconstructing the history of common marmosets (*Callithrix jacchus*) from high-quality, whole-genome, population-level data. Mol Biol Evol. 42(6):msaf119.

Spence JP, Song YS. 2019. Inference and analysis of population-specific fine-scale recombination maps across 26 diverse human populations. Sci Adv. 5(10):eaaw9206.

Spencer CC, Deloukas P, Hunt S, Mullikin J, Myers S, Silverman B, Donnelly P, Bentley D, McVean G. 2006. The influence of recombination on human genetic diversity. PLoS Genet. 2(9):e148.

Stapley J, Feulner PGD, Johnston SE, Santure AW, Smadja CM. 2017. Variation in recombination frequency and distribution across eukaryotes: patterns and processes. Philos Trans R Soc Lond B Biol Sci. 372(1736):20160455.

Stevison LS, Woerner AE, Kidd JM, Kelley JL, Veeramah KR, McManus KF, Great Ape Genome Project, Bustamante CD, Hammer MF, Wall JD. 2016. The time scale of recombination rate evolution in great apes. Mol Biol Evol. 33(4):928–945.

Stumpf MPH, McVean GAT. 2003. Estimating recombination rates from population-genetic data. Nat Rev Genet. 4(12):959–968.

Sweeney CG, Curran E, Westmoreland SV, Mansfield KG, Vallender EJ. 2012. Quantitative molecular assessment of chimerism across tissues in marmosets and tamarins. BMC Genomics. 13:98.

Tardif SD, Jaquish CE. 1997. Number of ovulations in the marmoset monkey (*Callithrix jacchus*): relation to body weight, age and repeatability. Am J Primatol. 42(4):323–329.

Tardif SD, Smucny DA, Abbott DH, Mansfield K, Schultz-Darken N, Yamamoto ME. 2003. Reproduction in captive common marmosets (*Callithrix jacchus*). Comp Med. 53(4):364–368.

Thomas GWC, Wang RJ, Puri A, Harris RA, Raveendran M, Hughes DST, Murali SC, Williams LE, Doddapaneni H, Muzny DM, et al. 2018. Reproductive longevity predicts mutation rates in primates. Curr Biol. 28(19):3193–3197.

van der Auwera G, O’Connor B. 2020. Genomics in the Cloud. 1st edition. O’Reilly Media.

Versoza CJ, Ehmke EE, Jensen JD, Pfeifer SP. 2025a. Characterizing the rates and patterns of *de novo* germline mutations in the aye-aye (*Daubentonia madagascariensis*). Mol Biol Evol. 42(3):msaf034.

Versoza CJ, Lloret-Villas A, Jensen JD, Pfeifer SP. 2025b. A pedigree-based map of crossovers and non-crossovers in aye-ayes (*Daubentonia madagascariensis*). Genome Biol Evol. 17(5):evaf072.

Versoza C, Weiss S, Johal R, La Rosa B, Jensen JD, Pfeifer SP. 2024. Novel insights into the landscape of crossover and non-crossover events in rhesus macaques (*Macaca mulatta*). Genome Biol Evol. 16(1):evad223.

Wall JD, Robinson JA, Cox LA. 2022. High-resolution estimates of crossover and noncrossover recombination from a captive baboon colony. Genome Biol Evol. 14(4): evac040.

Ward JM, Buslov AM, Vallender EJ. 2014. Twinning and survivorship of captive common marmosets (*Callithrix jacchus*) and cotton-top tamarins (*Saguinus oedipus*). J Am Assoc Lab Anim Sci. 53(1):7–11.

Williams AL, Patterson N, Glessner J, Hakonarson H, Reich D. 2012. Phasing of many thousands of genotyped samples. Am J Hum Genet. 91(2):238–251.

Wislocki GB. 1939. Observations on twinning in marmosets. Am J Anat. 64:445–483.

Xue C, Raveendran M, Harris RA, Fawcett GL, Liu X, White S, Dahdouli M, Rio Deiros D, Below JE, Salerno W, et al. 2016. The population genomics of rhesus macaques (*Macaca mulatta*) based on whole-genome sequences. Genome Res. 26(12):1651–1662.

Xue C, Rustagi N, Liu X, Raveendran M, Harris RA, Venkata MG, Rogers J, Yu F. 2020. Reduced meiotic recombination in rhesus macaques and the origin of the human recombination landscape. PLoS One. 15(8):e0236285.

Yang C, Zhou Y, Marcus S, Formenti G, Bergeron LA, Song Z, Bi X, Bergman J, Rousselle MMC, Zhou C, et al. 2021. Evolutionary and biomedical insights from a marmoset diploid genome assembly. Nature. 594(7862):227–233.

Zoonomia Consortium. 2020. A comparative genomics multitool for scientific discovery and conservation. Nature. 587(7833):240–245.

