## Supplementary Materials for "Inferring the landscapes of mutation and recombination in the common marmoset (*Callithrix jacchus*) in the presence of twinning and hematopoietic chimerism"

**Supplementary Table 1. Summary of variant and invariant datasets**. A total of 7.2 million autosomal single nucleotide polymorphisms (SNPs) with a transition-transversion ratio (Ts/Tv) of 2.2 were discovered in the accessible genome of the 15 individuals included in this study.

| chr | length | # SNPs | # invariant sites | Ts/Tv |
|---|---|---|---|---|
| 1 | 216,975,769 | 524,830 | 92,335,456 | 2.21 |
| 2 | 202,808,461 | 497,309 | 83,856,609 | 2.14 |
| 3 | 189,455,076 | 380,256 | 61,918,066 | 2.07 |
| 4 | 173,414,057 | 460,590 | 74,586,824 | 2.09 |
| 5 | 161,716,902 | 427,874 | 97,377,364 | 2.20 |
| 6 | 159,674,559 | 412,940 | 65,347,930 | 2.17 |
| 7 | 156,129,252 | 479,974 | 84,619,705 | 2.21 |
| 8 | 126,104,592 | 344,488 | 52,405,398 | 2.13 |
| 9 | 132,900,640 | 327,124 | 61,796,278 | 2.15 |
| 10 | 136,971,485 | 421,159 | 66,986,906 | 2.15 |
| 11 | 128,529,401 | 368,614 | 60,643,965 | 2.17 |
| 12 | 123,697,827 | 373,566 | 65,153,373 | 2.23 |
| 13 | 117,638,787 | 274,234 | 54,984,316 | 2.13 |
| 14 | 112,969,529 | 289,729 | 53,477,644 | 2.11 |
| 15 | 98,621,277 | 305,926 | 47,108,341 | 2.17 |
| 16 | 98,110,366 | 241,857 | 38,432,144 | 2.05 |
| 17 | 74,700,100 | 194,268 | 30,264,449 | 2.11 |
| 18 | 47,063,576 | 140,577 | 23,830,279 | 2.14 |
| 19 | 50,540,917 | 214,633 | 29,637,108 | 2.19 |
| 20 | 44,412,365 | 144,326 | 24,251,532 | 2.26 |
| 21 | 50,614,742 | 154,261 | 18,311,224 | 2.11 |
| 22 | 49,900,457 | 219,893 | 27,282,321 | 2.27 |
| ∑ or ∅ | **2,652,950,137** | **7,198,428** | **1,214,607,232** | **2.16** |

**Supplementary Table 2. Divergence time estimates.** Comparison of inferred common marmoset divergence times based on the observed mean neutral divergence rate of 0.0029 between *C. jacchus* and *C. kuhlii* for two possible generation times (1.5 years and 2.0 years; Tardif et al. 2003; Okano et al. 2012; Schultz-Darken et al. 2016; Han et al. 2022) and for pedigree-based mutation rates previously estimated in common marmosets ($0.43 \times 10^{-8}$ mutations per base pair per generation; Yang et al. 2021) as well as in the closely related (but non-chimeric) owl monkeys ($0.81 \times 10^{-8}$ mutations per base pair per generation; Thomas et al. 2018).

| | | **pedigree-based mutation rate** | |
|---|---|---|---|
| | | **4.30E-09** | **8.10E-09** |
| **generation time** | **1.5 years** | 0.34 mya | 0.18 mya |
| | **2.0 years** | 0.49 mya | 0.24 mya |

#### Supplementary Figure 1

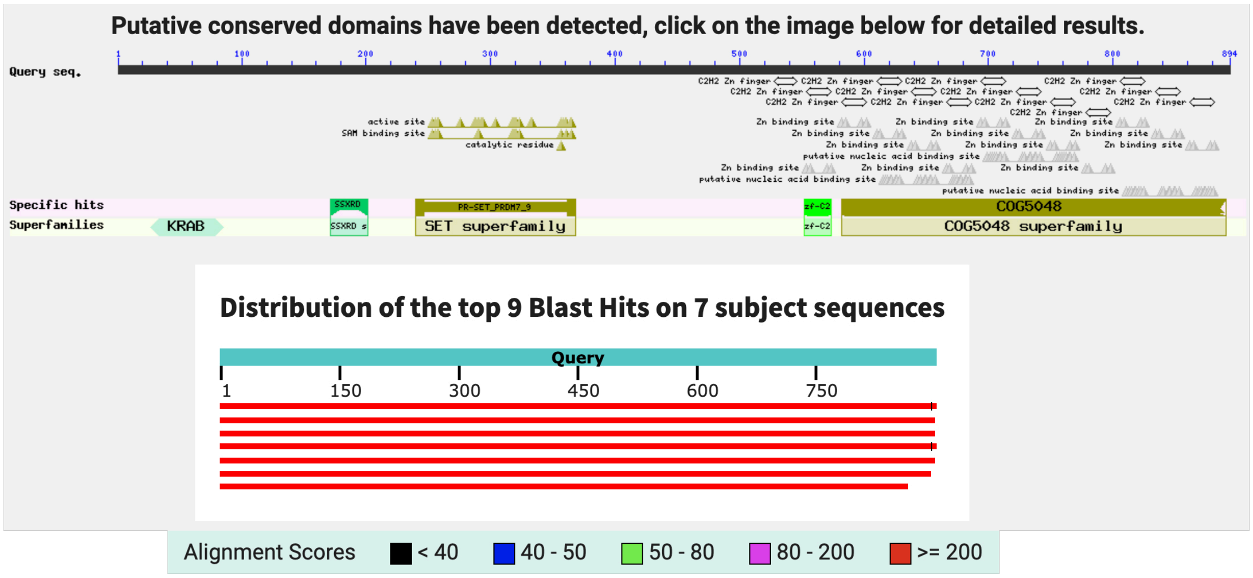


**Supplementary Figure 1.** Putatively conserved domains (including the a Krueppel-associated box [KRAB] domain, the SSX repressor domain [SSXRD], and the PR/SET domain) and the C2H2-type zinc finger array detected in the protein-protein BLAST search of the marmoset PRDM9 amino acid sequence against the NCBI non-redundant protein sequences database.

#### Supplementary Figure 2

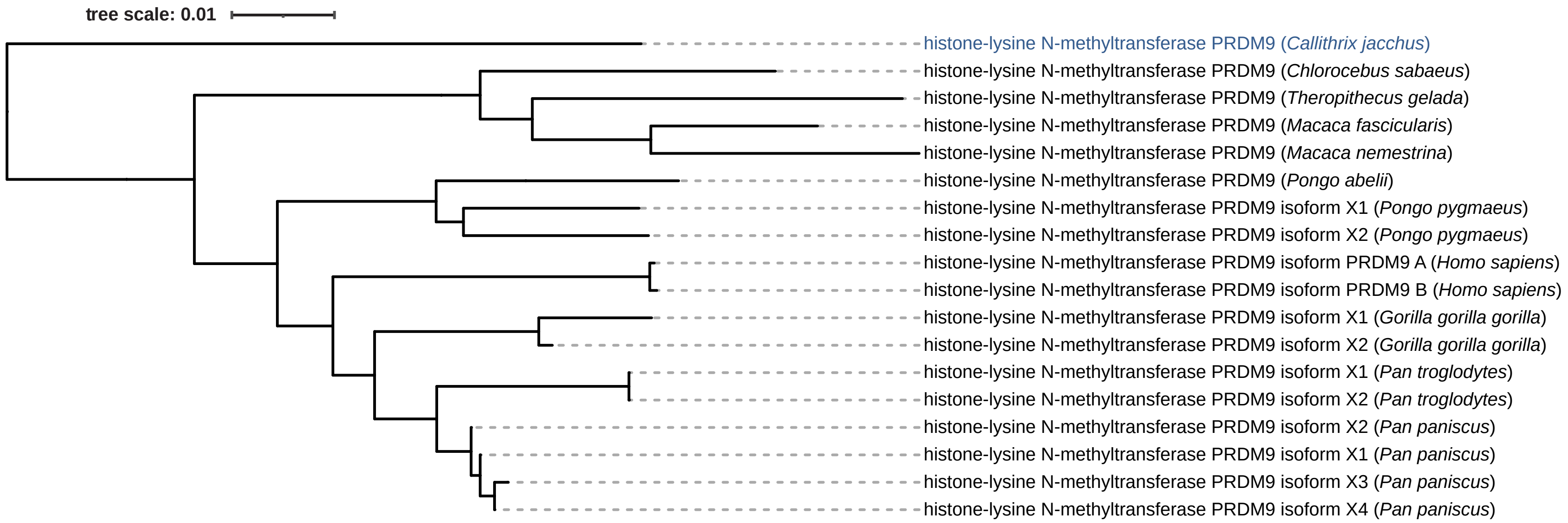


**Supplementary Figure 2.** Phylogenetic tree resulting from the protein-protein BLAST search of the marmoset (*C. jacchus*) PRDM9 amino acid sequence (shown in blue) against the NCBI non-redundant protein sequences database.

#### Supplementary Figure 3

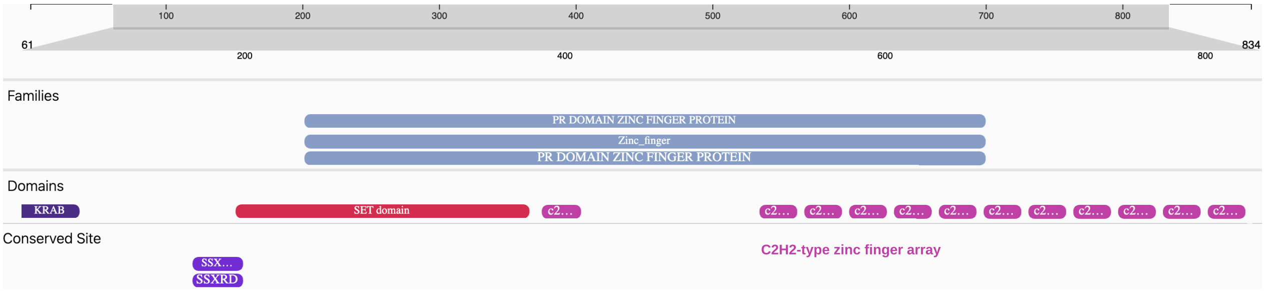


**Supplementary Figure 3.** Protein domains within the marmoset (*C. jacchus*) PRDM9 amino acid sequence as predicted by the InterPro web server.

**Supplementary Figure 4**

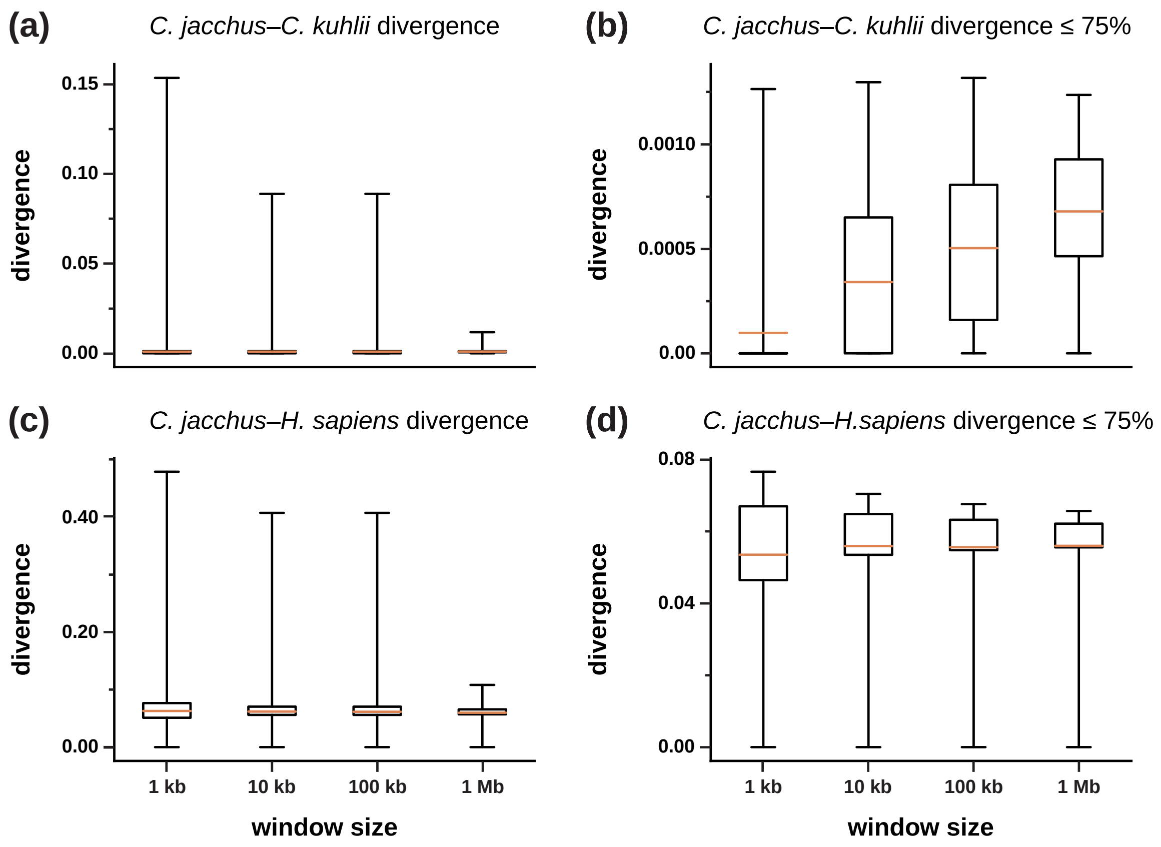


**Supplementary Figure 4.** Distributions of neutral divergence. Top: distribution of neutral divergence between *C. jacchus* and *C. kuhlii* for (a) all 1 kb, 10 kb, 100 kb and 1 Mb genomic windows and (b) for windows in the lower three quartiles. Bottom: distribution of neutral divergence between *C. jacchus* and *H. sapiens* for (c) all 1 kb, 10 kb, 100 kb and 1 Mb genomic windows and (d) for windows in the lower three quartiles. Orange bars represent the median neutral divergence, boxes represent the first and third quartiles, and whiskers represent the minimum and maximum values.

### Supplementary Figure 5

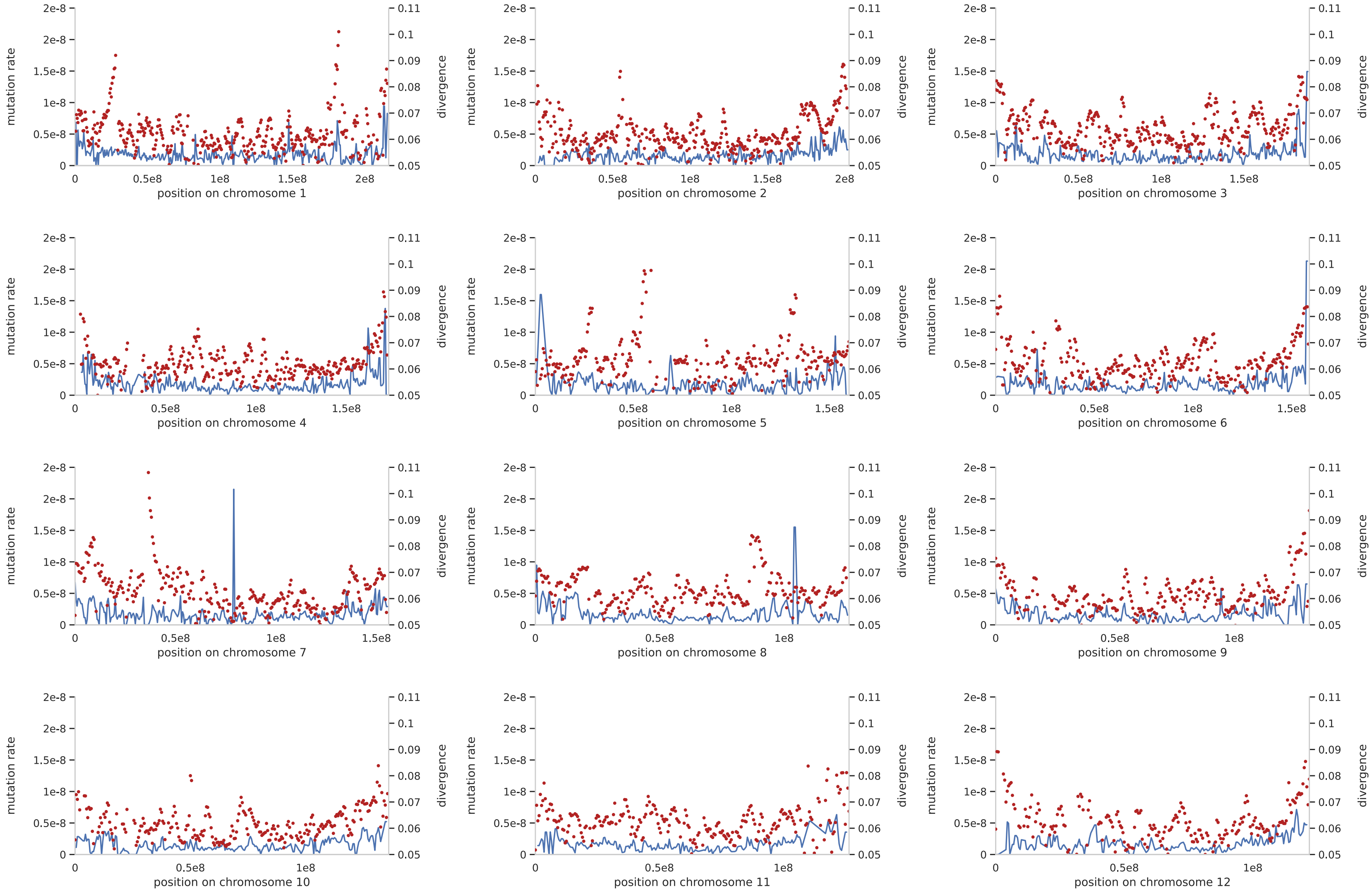

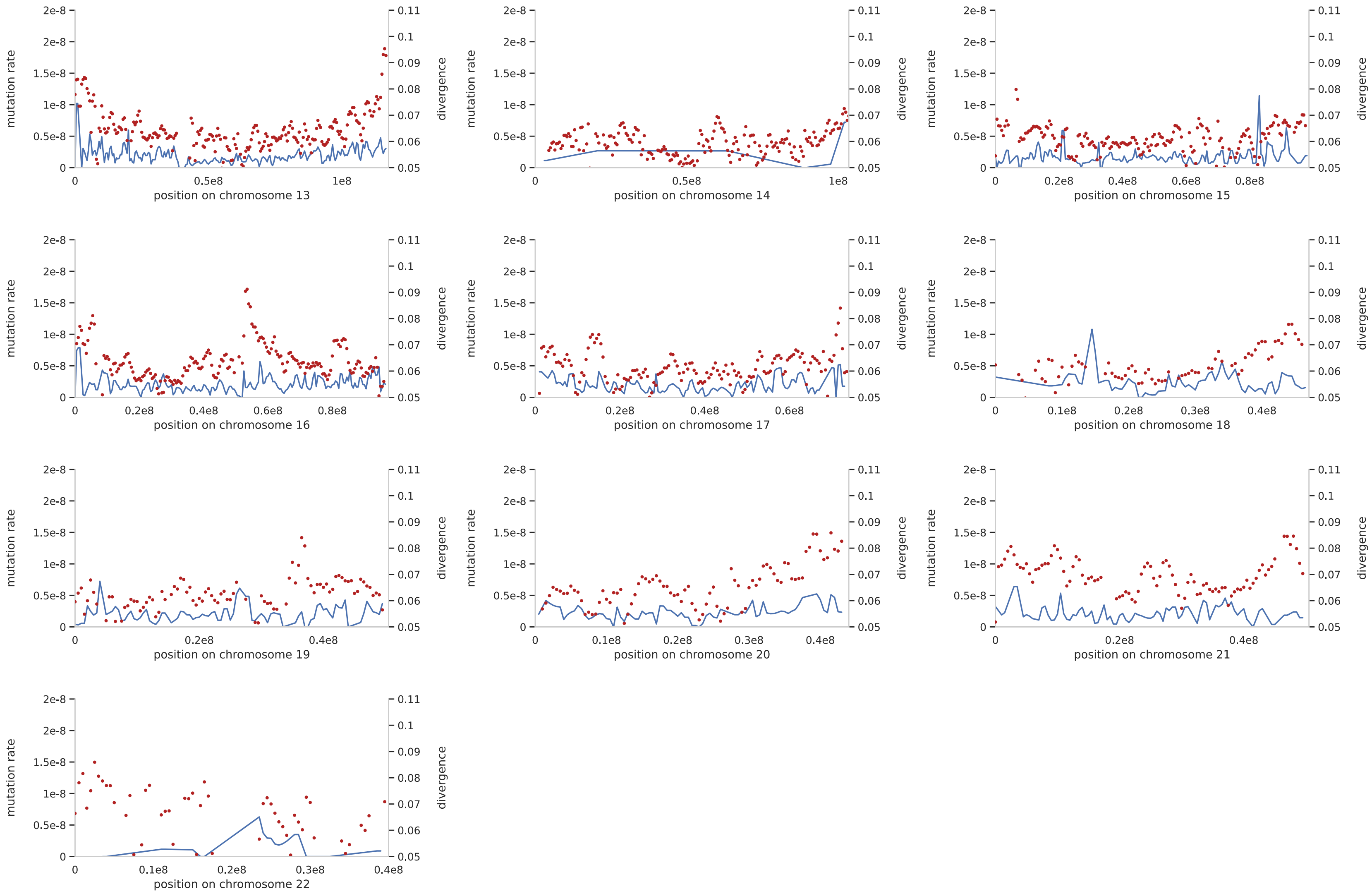


**Supplementary Figure 5.** Fine-scale per-site per-generation neutral mutation rates indirectly obtained from the *C. jacchus*–*C. kuhlii* divergence (shown in blue on the left y-axis) and neutral divergence rate estimates based on *C. jacchus*–*H. sapiens* alignments (shown in red on the right y-axis) along each autosome for genomic windows of size 1 Mb, with a 500 kb step size.

**Supplementary Figure 6**

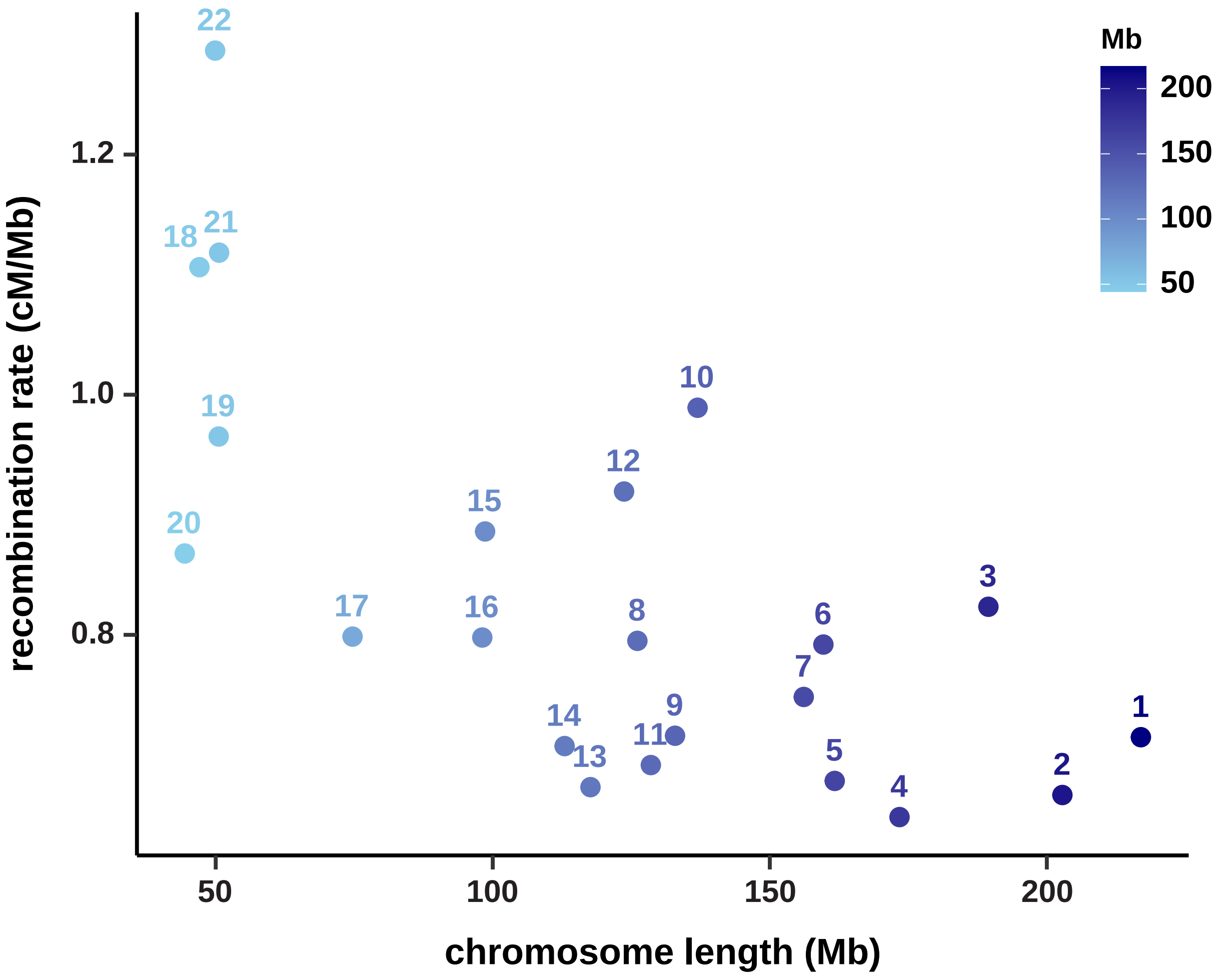


**Supplementary Figure 6.** Relationship between the per-site per-generation recombination rates as inferred by LDhat and the size of each autosome (chromosomes 1-22).
This plot was produced using a script provided by Bascón-Cardozo et al. 2024 (https://github.com/Karenbc/Recombination-rates-and-genomic-features-Blackcap).

### Supplementary Figure 7

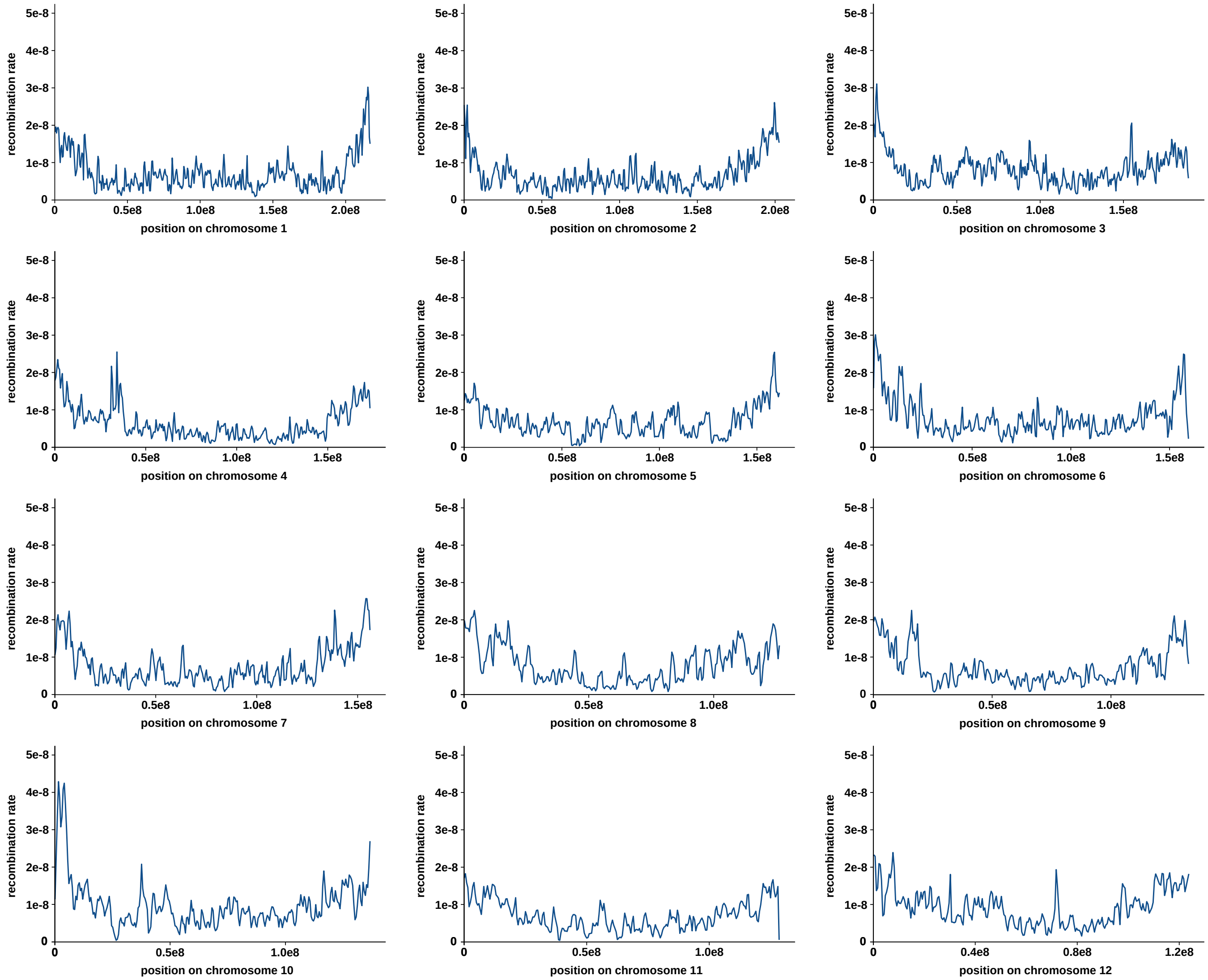

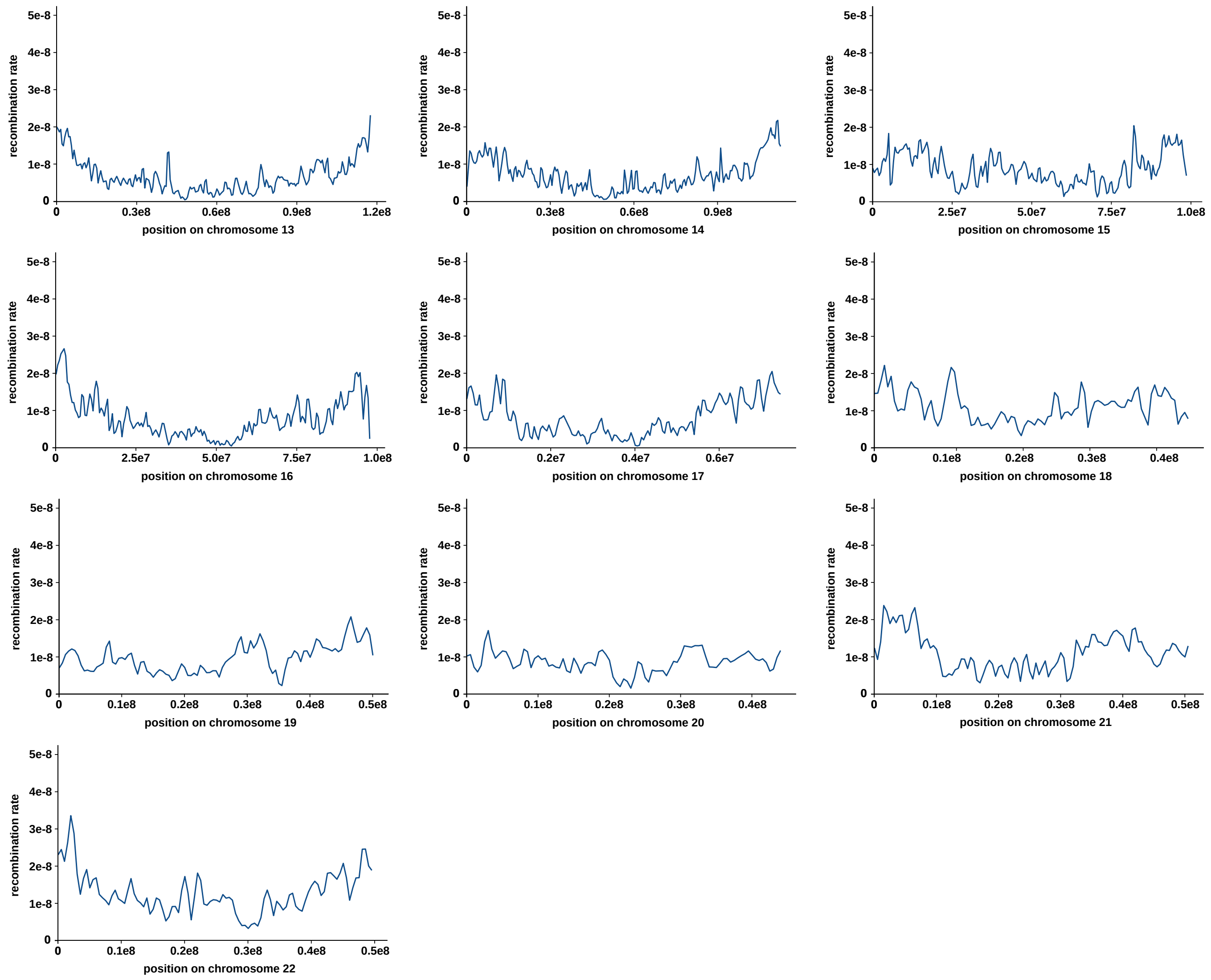


**Supplementary Figure 7.** Fine-scale per base pair per generation recombination rates as inferred by LDhat for genomic windows of size 1 Mb, with a 500 kb step size.

**Supplementary Figure 8**

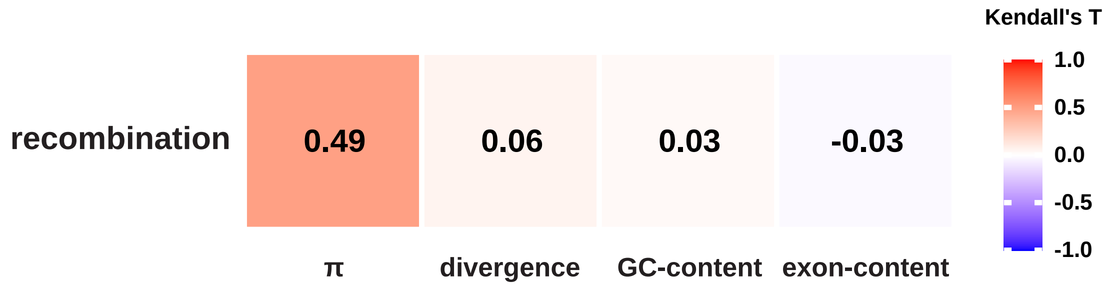


**Supplementary Figure 8.** Relationship between the per-site per-generation recombination rates as inferred by LDhat and four genomic features: nucleotide diversity ($\pi$), divergence, GC-content, and exon-content. Partial Kendall's rank correlations were calculated in 100 kb-windows in which at least 50% of sites were accessible.

**Supplementary Figure 9**

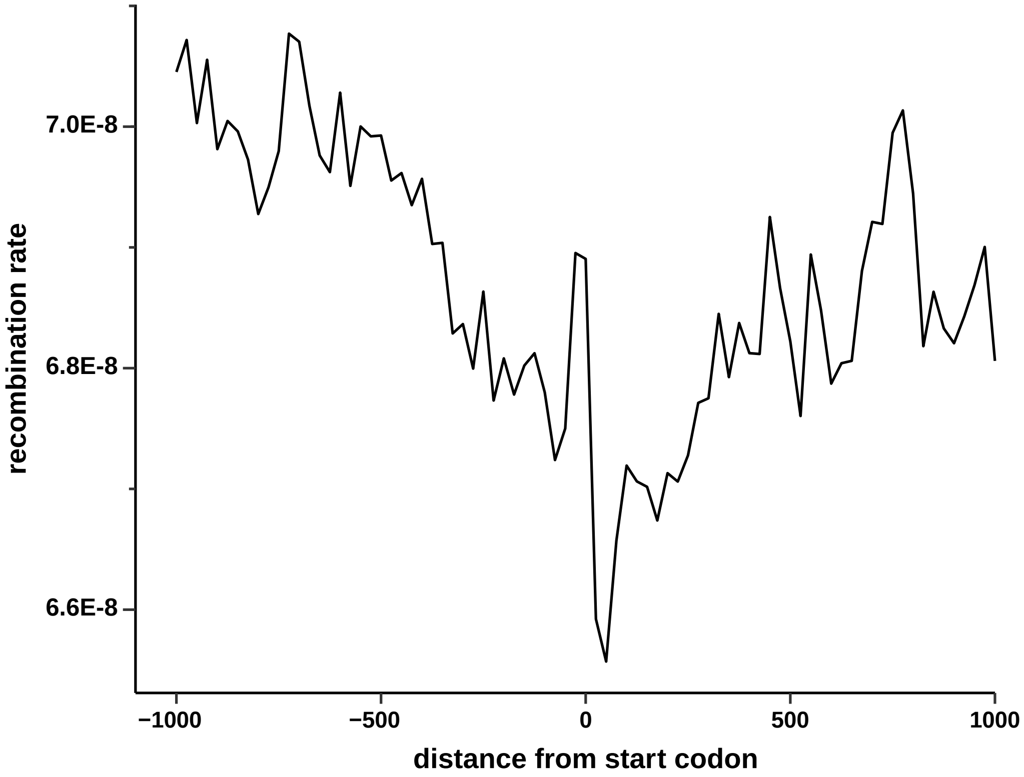


**Supplementary Figure 9.** Per-site per-generation recombination rates as inferred by LDhat as a function of distance to the nearest start codon (averaged across 1 kb windows).

**Supplementary Figure 10**

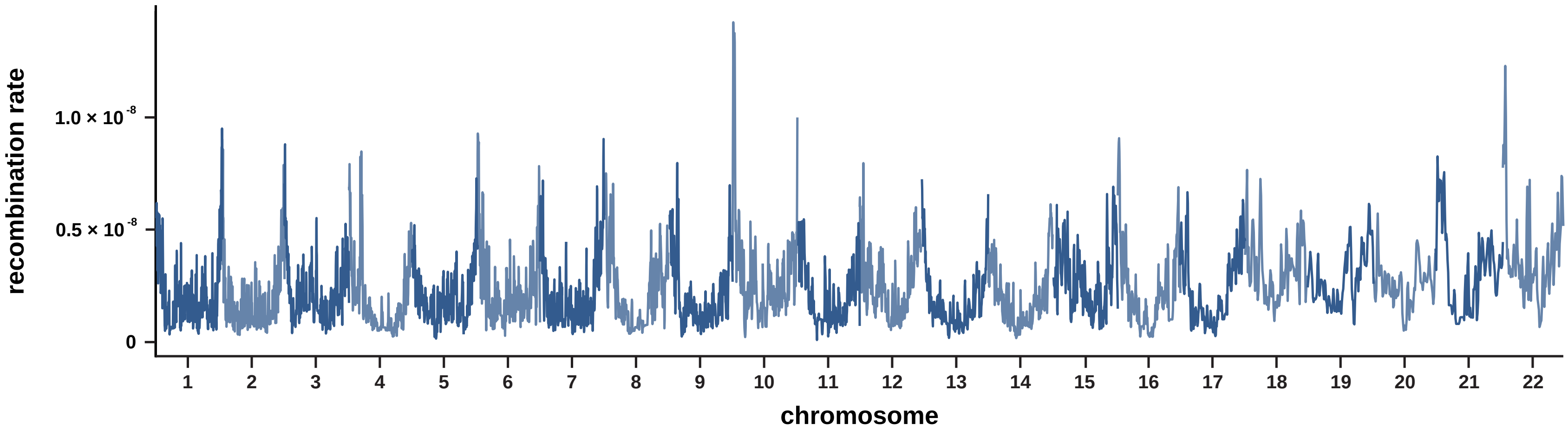


**Supplementary Figure 10.** Fine-scale per base pair per generation recombination rates as inferred by pyrho along each autosome.
